# Far-Red Light Photoacclimation Rewires Antenna-Photosystem Organization in Cyanobacterial Cells

**DOI:** 10.64898/2026.10.21.760696

**Authors:** Sofie van Dorst, Vera M. Selinger, Dennis J. Nürnberg, Wojciech Wietrzynski, Benjamin D. Engel

**Affiliations:** Biozentrum, University of Basel, Spitalstrasse 41, 4056 Basel, Switzerland; Institute for Experimental Physics, Freie Universität Berlin, Arnimallee 14, 14195 Berlin, Germany; Institute for Biology/Microbiology, Martin-Luther-Universität Halle-Wittenberg, Kurt-Mothes-Straße 3, 06120 Halle (Saale), Germany

## Abstract

Thylakoid membrane remodeling regulates photosynthetic activity in response to changing light, but it is unknown whether such membrane-scale control exists in cyanobacteria, which lack the spatially segregated membrane domains of green algae and plants. Using cryo-electron tomography guided by single-cell hyperspectral correlative imaging, we link spectral acclimation state to thylakoid architecture in far-red light adapting cyanobacteria. We determine the native structures of phycobilisome-photosystem II supercomplexes at subnanometer resolution and show that light-dependent molecular changes propagate across scales to restructure the photosynthetic machinery throughout the thylakoids. We further reveal persistent membrane heterogeneity after prolonged far-red light acclimation and identify a low abundance phycobilisome-photosystem I supercomplex in far-red light conditions. Together, our results frame cyanobacterial thylakoids as plastic membranes that remodel antenna-photosystem structure and organization to regulate photosynthesis.

## Introduction

Photosynthesis was long thought to be constrained by a strict red-limit at 700 nm, because the absorption of light by the near-universal photosynthetic pigment chlorophyll (Chl) *a* drops steeply beyond this wavelength, and these photons do not provide sufficient energy to drive this process^1^. However, the discovery of red-shifted chlorophylls Chl *d* ^2^ and Chl *f* ^3^, and a conserved Far-Red Light Photoacclimation (FaRLiP) gene cluster^4^, have shown that photosynthesis can expand into much more restrictive light regimes. The FaRLiP gene cluster encodes for red-shifted paralogs of ∼20 genes of the photosynthetic apparatus, including most components of photosystem I and II (PSI and PSII), the light harvesting phycobilisomes (PBS), and regulatory genes, enabling a select group of cyanobacteria to sustain photosynthesis with far-red photons >700 nm in wavelength^5,6^. This adaptation is rare, but phylogenetically widespread, and is thought to trace back to early days of oxygenic photosynthesis^6,7^. FaRLiP allows these cyanobacteria to thrive in light-limited and competitive niches where far-red wavelengths dominate, such as dense microbial mats, stromatolites, and endolithic communities^8–10^. The principle of expanding photosynthesis into wavelengths beyond those used by Chl *a* has motivated efforts to engineer red-shifted photosynthesis in plants to improve canopy light use and crop productivity^11,12^. However, a lack of understanding of the molecular mechanisms in cyanobacteria remains a major bottleneck.

FaRLiP cyanobacteria dynamically switch between white light (WL, *wl*) photosynthetic machinery, expressed in visible light regimes (400-700 nm), and far-red light (FRL, *frl)* machinery, expressed upon exclusive exposure to FRL wavelengths^13^. The transition to far-red photosynthesis is slow, lasting days to weeks, which can be explained by the significant investments into production of far-red absorbing chlorophylls and their incorporation into new photosystems, all under the initially light-limiting condition^5,14^. Genetic, biochemical, and structural evidence show that FaRLiP-associated changes in photosynthetic membranes range in scale from new photosystem subunits and remodeling of PBS to changes to the overall thylakoid network^15–19^. However, conclusive evidence is lacking for the structure of the *frl*-PBS inside cells, its mode of association to PSI and PSII, and the FRL-induced reorganization of photosynthetic protein complexes in the larger context of the thylakoid membranes. The inducible wholesale replacement of photosynthetic machinery with structurally distinct far-red isoforms makes the FaRLiP process exceptionally suited for probing how cyanobacteria coordinate photosynthetic remodeling from the molecular to membrane scale.

Cryogenic electron tomography (cryo-ET) has proven to be a powerful method for analyzing the molecular architecture of photosynthetic membranes inside the native environment of the cell^20,21^. By combining cryo-ET with a novel hyperspectral correlative light and electron microscopy (hsCLEM) workflow, we connect high-resolution protein structure, native membrane architecture, and photosynthetic physiology to reveal how *wl*-, and *frl*-photosynthetic complexes are organized within thylakoid membranes of the cyanobacterium *Chroococcidiopsis thermalis* PCC7201.

## Results and Discussion

### FRL induces remodeling of PBS structure and thylakoid organization in C. thermalis

*C. thermalis* cells grown in WL displayed concentric layers of curved thylakoids, typical of coccoid cyanobacteria^22^ (Fig. 1A; Fig. S2A-F). The cytoplasmic surface of the thylakoids was extensively decorated with tightly packed rows of large hemidiscoidal *wl*-PBS, which stoichiometrically associated with linear arrays of PSII protruding from the luminal surface of the membrane (Fig. 1A,C). Inter-thylakoid membrane distance was approximately spaced to the dimensions of the *wl*-PBS (69 ± 20 nm) (Fig. S3C), and glycogen granules were frequently associated with the thylakoids (Fig. 1A,C).

**Figure 1.**
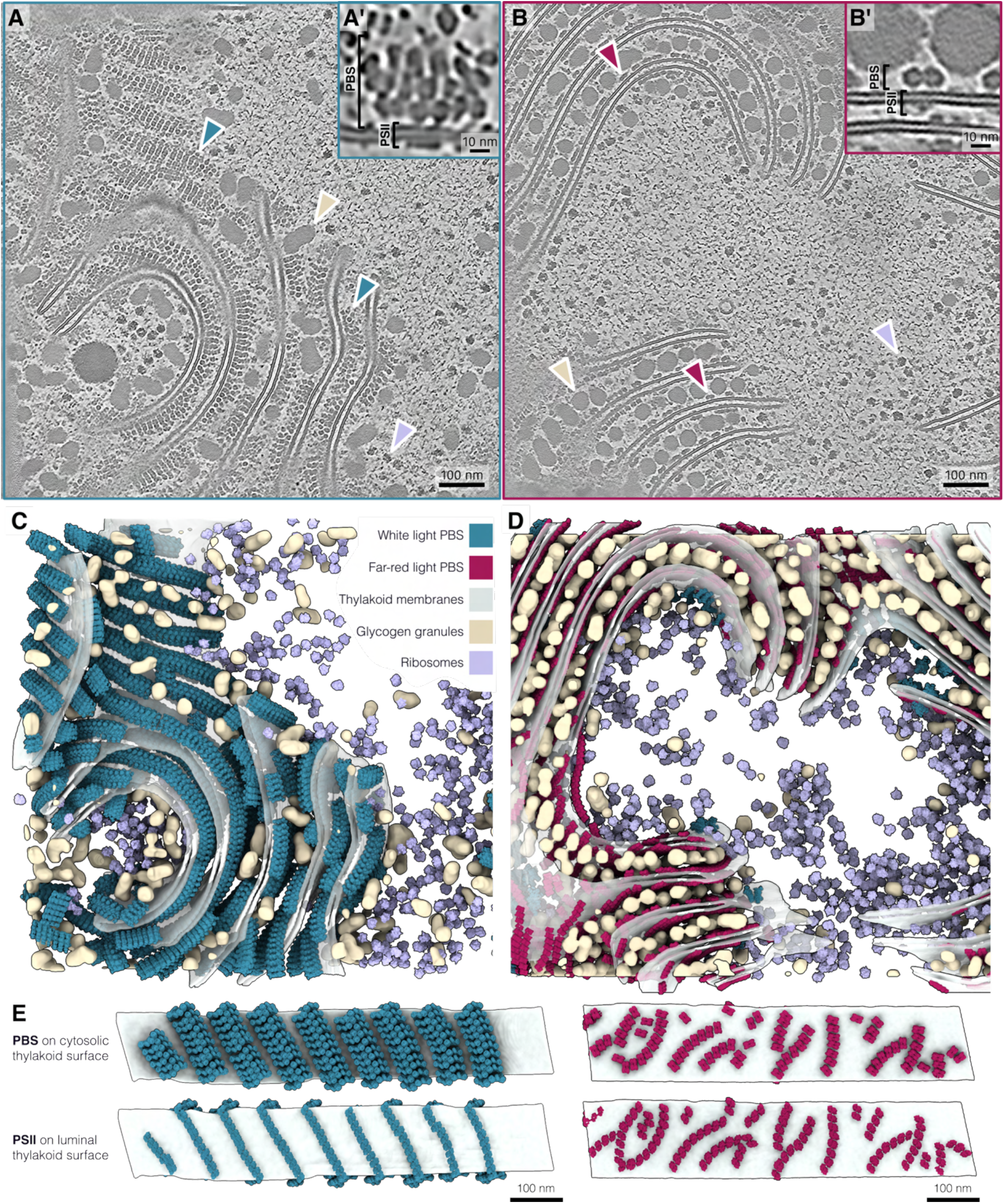
FRL induces remodeling of PBS structure and organization in *C. thermalis*. **A, B**, Representative tomographic slices of *C. thermalis* cells acclimated to white light (WL, **A**) or far-red light (FRL, **B**). WL cells display thylakoids decorated with rows of hemidiscoidal *wl*-PBS associated with PSII, while FRL cells contain bicylindrical *frl*-PBS associated with PSII. Arrowheads indicate *wl*-PBS in blue, *frl*-PBS in red, glycogen granules in tan, and ribosomes in lilac. **A′, B′**, Zoomed-in views of PBS-PSII complexes in WL and FRL cells, respectively. PBS are located on the cytosolic side of the membrane, while PSII spans the membrane and protrudes from the luminal side. **C, D**, Segmentations and map-backs of subtomogram averages corresponding to the tomograms of the WL cell in **A** and the FRL cell in **B**. *wl*-PBS are shown in blue, *frl*-PBS in red, thylakoid membranes in light grey, glycogen granules in tan, and ribosomes in lilac. **E**, Segmented sections of thylakoid membranes from WL and FRL tomograms, shown with PBS map-backs on the cytosolic surface and corresponding PSII map-backs visible on the luminal surface. The reduced size of *frl-*PBS allows denser packing of PBS-PSII supercomplex rows. Scale bars 100 nm in **A**, **B** and **E;** 10 nm in **A′** and **B′**.

Acclimation of *C. thermalis* to FRL led to striking remodeling of the PBS. Consistent with previous work on isolated *frl*-PBS^15,23,24^, the dominant population of PBS in FRL was a reduced, bicylindrical structure lacking rods, thus resulting in threefold smaller lateral footprint of the PBS (width of ∼22 nm versus ∼62 nm for *wl*-PBS) (Fig. 1E; S6A,D). These *frl*-PBS formed rows similar to *wl*-PBS, both with a center-to-center spacing of ∼14 nm. However, the smaller *frl*-PBS structure permits a higher density of *frl*-PBS-PSII supercomplexes in the same membrane area while also occupying less cytosolic space above the membrane than in WL (Fig. 1E). Therefore, while the overall architecture of thylakoid membranes is preserved in FRL, the reduction in PBS size allows thylakoid membranes to pack closer together (47 ± 23 nm) (Fig. S3B,C), as described previously^16^.

PBS remodeling caused substantial repercussions on the overall molecular organization of the thylakoids. In WL, the hemidiscoidal *wl-*PBS covered much of the cytosolic thylakoid surface in flat and convex regions, whereas they were sparse in concave regions, possibly due to steric hindrance of the large *wl-*PBS arrays (Fig. S6B,C). A closer look at these membranes revealed that they were also decorated with small densities protruding on the cytosolic side (Fig. 2). The size, organization, and abundance of these densities are consistent with the cytosolic domain of PSI. In WL, there were few PSI in *wl-*PBS-rich regions (Fig. 2B,E,H), whereas PSI was abundant in the concave membranes where *wl-*PBS were sparse (Fig. 2B,C,F,I). These results support the presence of two distinct and spatially separated membrane domains in WL: one enriched in PBS-PSII supercomplexes and the other enriched in PSI^25^. By contrast, FRL thylakoids were decorated on both concave and convex surfaces by rows of *frl-*PBS, resulting in global intermixing of PBS-PSII supercomplexes with mostly trimeric PSI densities (Fig. 2D,G,J). Consequently, there is no obvious partitioning of thylakoid membranes into separate domains in FRL.

**Figure 2.**
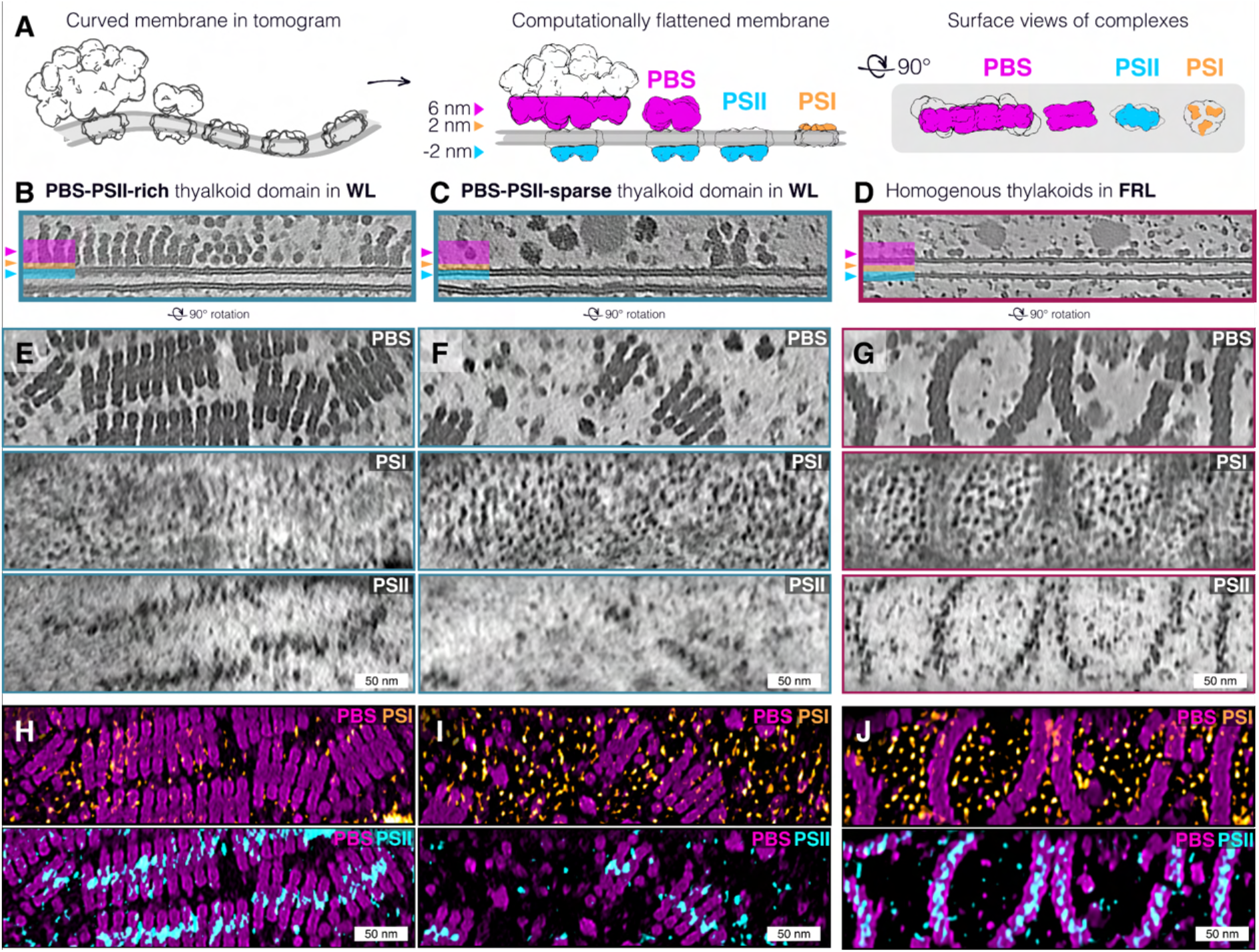
Light-dependent domain organization of photosynthetic complexes in native thylakoid membranes. **A**, Schematic representation of how 3D segmentations of curved membranes in a tomogram are computationally flattened. Visualization of tomographic density at various distances from the membrane surface (surface views, sometimes called “membranograms”) reveals densities corresponding to the various photosynthetic complexes: Densities are shown at ∼6 nm on the cytosolic side, corresponding primarily to PBS, but also including other cytosolic densities such as ATP synthase and ribosomes; ∼2 nm on the cytosolic side, corresponding mostly to PSI; and ∼2 nm on the luminal side, corresponding mostly to PSII. **B-D** Flattened membranes from WL and FRL tomograms, perpendicular view. WL thylakoids show distinct PBS–PSII-rich regions (**B**) and PBS–PSII sparse regions (**C**), whereas FRL thylakoids display a more homogeneous distribution of photosynthetic complexes (**D**). **E–G**, Surface views of membranes from B-D, summed over 2–3 slices at defined distances from the thylakoid membrane, showing densities corresponding to PBS (top), PSI (middle), and PSII (bottom). **H–J**, Overlays of surface-view densities of WL PBS–PSII-rich region (**H**), WL PBS–PSII-sparse region (**I**), and FRL region (**J**). PBS are shown in magenta, PSI in orange, and PSII in blue. Scale bars 50 nm.

PBS-PSII rows thus appear to be a determinant factor in the mesoscale organization of cyanobacterial thylakoid membranes. By changing from bulkier, potentially more geometrically constrained rows in WL into narrower, potentially more flexible rows in FRL, thylakoids dynamically alter the distribution and organization of their PSI and PSII complexes. This indicates that despite the lack of clear lateral heterogeneity of photosystems enforced by thylakoid stacking, as found in green algae^26^ and plants^27^, cyanobacteria can form and dissolve discrete thylakoid membrane domains by altering the molecular architecture of their photosynthetic machinery.

### Alternative arrangement of PBS-PSII supercomplexes under FRL conditions

Subtomogram averaging (STA) of *wl-*PBS-PSII supercomplexes inside WL-grown *C. thermalis* cells revealed a hemidiscoidal PBS with a pentacylindrical allophycocyanin core and phycocyanin rods protruding radially from the core. This *wl-*PBS closely resembles the structure of the pentacylindrical PBS isolated from *Anabaena* sp. PCC 7120^28^ (Fig. 3A,C; Fig. S8M,N). Minor repositioning of the peripheral rods Rs2 and Rs2’ compared to the isolated structure was necessary to accommodate the interdigitated packing of *wl-*PBS into their rows (Fig. S8L-N). The peripheral rods are less well resolved in the STA map, suggesting that they may be more flexible or heterogeneous (Fig. 3C). The PBS associates with PSII in an overall arrangement similar to the tricylindrical PBS-PSII supercomplex in the cyanobacterium *Arthrospira* sp. FACHB-439^21^, where each PBS docks asymmetrically onto the interface of two PSII dimers in a row, but with a minor repositioning of the exact docking site (Fig. 3F; S8A-C). Our *wl-*PSII average is also in excellent agreement with the *Arthrospira* structure, but altogether lacks the Lpp1 peptide described in *Arthrospira*^21^ and the red alga *Porphoridium purpureum*^29^ as a PBS-PSII linker protein (Fig. S8H). Moreover, we noted that approximately half of the row-capping *wl*-PBS possessed a repositioned toroid that extends from the Rs1 position into the open space (here, we designate this position Rs1*) (Fig. S11).

**Figure 3.**
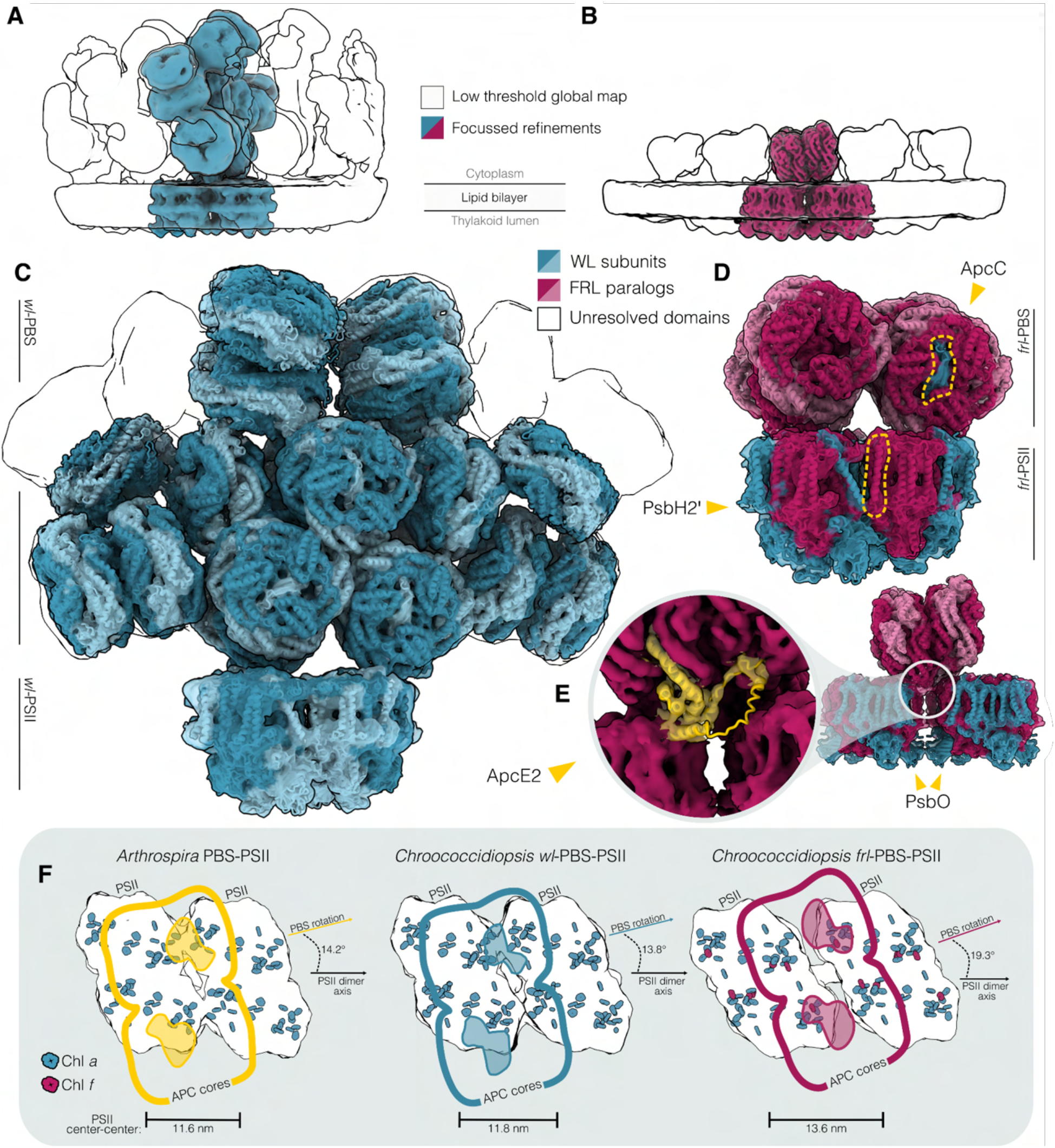
Molecular details of the PBS-PSII supercomplex in WL and FRL conditions. **A, B**, Side views of subtomogram averages of the *wl*-PBS-PSII (**A**) and *frl*-PBS-PSII (**B**) supercomplexes in their row formation. Focused refinements of one PBS and two associated PSII dimers are shown in blue for WL and red for FRL, overlaid with a low-threshold, locally filtered map showing the broader row organization as an outline. In both conditions, a PBS binds at the interface of neighboring PSII dimers, forming extended PBS-PSII arrays. In WL, the PBS docks asymmetrically and is positioned mainly above one PSII dimer, whereas in FRL, the PBS adopts a more symmetric position between two PSII dimers. **C**, Front view of the *wl*-PBS-PSII subtomogram average (Global resolution: 9.1 Å, PBS resolution: 7.2 Å, PSII resolution: 9.9 Å, **Fig. S7J-K**) fitted with molecular models. The *wl*-PBS closely resembles the pentacylindrical PBS from *Anabaena* sp. PCC 7120 (PDB: 7EYD), while PSII was fitted with the *Thermosynechococcus vulcanus* PSII structure (PDB: 8IR7). Peripheral PBS rods are less well resolved, owing to their flexibility or heterogeneity, and are shown as outlines. **D**, Front view of the bicylindrical *frl*-PBS-PSII subtomogram average (Global resolution: 10.63 Å, PBS resolution: 8.2 Å, PSII resolution: 8.9 Å, **Fig. S7L-M**) fitted with structures of the *C. thermalis frl*-PBS (PDB: 9I1R) and *frl*-PSII (PDB: 9T5T). FaRLiP-encoded components are shown in red, while WL-conserved components are shown in blue. ApcC, present as a single copy per APC core, and the FRL-specific PSII subunit PsbH2′ are highlighted. **E**, Density at the *frl*-PBS-PSII interface shows the ApcE2 extension contacting *frl*-PSII. The corresponding full-length ApcE2 AlphaFold prediction is fitted into the map density and shown in yellow. **F**, Comparison of PBS-PSII docking geometries in PBS-PSII from *Arthrospira* sp. FACHB-439 (PDB: 8WQL), *C. thermalis wl*-PBS-PSII, and *C. thermalis frl*-PBS-PSII. Two neighboring PSII dimers are aligned along their center-to-center axis, with PSII dimer spacing indicated. PBS positions are shown as colored outlines, and PBS–PSII contact sites are marked in color. Aromatic rings of PSII chlorophylls are shown with Chl *a* in blue and Chl *f* in red. While the *wl*-PBS-PSII geometry in *C. thermalis* resembles the asymmetric arrangement described in *Arthrospira*, *frl*-PBS-PSII is substantially rearranged, placing the PBS more centrally between neighboring PSII dimers and positioning its terminal bilins closer to Chl *f* sites.

STA in FRL-adapted cells revealed that the organization of the *frl-*PBS-PSII supercomplex is markedly different. As reported previously for an isolated *frl*-PBS complex^15^, this light-harvesting antenna consists of two allophycocyanin (APC) cores, each built of four toroid-shaped phycobiliprotein hexamers (Fig. 3B,D). It contains only a single copy of ApcC per APC core, in contrast with the apical and distal copy of ApcC present in each core of the *wl*-PBS (Fig. 3D). In both *wl-* and *frl-*supercomplexes, we observe no density corresponding to the PBS-PSII linker protein Lpp2 that was previously found in *Arthrospira* PBS-PSII (Fig. S8D-F,J-K). Our STA map of native *frl-*PSII at subnanometer resolution is in full agreement with the isolated single particle structure from *C. thermalis*^31^, including the recently identified subunit PsbH2’ uniquely found in *frl-*PSII (Fig. 3D).

Unlike the *wl-*PBS-PSII supercomplex, *frl-*PBS docks symmetrically onto its two associated PSII dimers, resulting in a notably larger spacing between neighboring PSII complexes as well as different PBS-PSII interaction sites (Fig. 3E,F), mediated by the FaRLiP-cluster-encoded ApcE2 protein (Fig. 3E). Compared to *wl-*ApcE, *frl-*ApcE (ApcE2) carries an extended, divergent loop (residues 110-150) that directly contacts PSII and appears to define the docking interaction of *frl*-PBS on PSII (Fig. S9). In our STA map, this loop approaches FRL-specific PSII subunits PsbH2 and D2 (PsbD2), ruling out PsbH2’ as the PBS-binding subunit, as has been proposed previously^31^ (Fig. 3E). This geometry positions the terminal bilin C201 such that the closest Chl *f* acceptor becomes C507 of *frl-*CP43 (PsbC2), with an edge-to-edge distance of 28.9 Å (Fig. 3F; Fig. S10). This favors the energy transfer route through Chl *f* C507 to the *frl*-PSII reaction center, which has been modeled to be an order of magnitude faster than the route involving the next closest Chl *f* site in CP47 (PsbB2)^32^ (Fig. S10). This supercomplex geometry explains why FRL energy transfer was measured faster *in vivo* when exciting through the *frl-*PBS, compared to the transfer in *frl-*PSII alone when exciting chlorophyll, despite the attached PBS creating a larger functional antenna size that should normally increase fluorescence lifetime^32^. Repositioning of the *frl-*PBS to channel excitation energy into CP43 increases the probability of energy transfer through this faster route, thereby enhancing the overall efficiency of the complex. The FRL-specific PBS-PSII organization thus directly contributes to faster energy transfer rates, a yield optimization strategy under energy-limited light conditions.

How exactly the difference in PBS-PSII row organization arises under FRL remains to be determined. The lack of protein density between neighboring PSII dimers in the row implies they interact in the membrane through lipids, while the luminal contacts appear to be mediated mainly by PsbO (Fig. 3E), which is not FRL-specific in *C. thermalis*. All *in situ* data suggest that PBS binding requires two neighboring PSII dimers^21,29^, implicating the *frl-*PBS in directing the altered supercomplex geometry. This remodeling therefore likely requires spatiotemporal coordination of *frl-*PBS and *frl-*PSII synthesis and assembly.

### Membrane organization of PSI and a rare *frl*-PBS-PSI supercomplex

In both WL- and FRL-grown *C. thermalis* cells, PSI densities were discernable directly adjacent to rows of PBS-PSII (Fig. 2H-J), similar to previous atomic force microscopy observations of isolated cyanobacterial membranes^33^ but with more variable orientations relative to the PBS-PSII supercomplexes. A large body of spectroscopic evidence suggests connectivity between PSII and PSI in cyanobacteria^34,35^. The close proximity of PSI to PBS-PSII arrays in our data points to the possibility of energy transfer from PSII to PSI.

Additionally, in cells acclimated to FRL, we observed rare solitary *frl*-PBS complexes at the membrane surface that lacked obvious luminal PSII densities (Fig. 4A,B). STA of these PBS positions revealed that this small subset of all identified PBS (4%) corresponds to a novel *frl*-PBS-PSI supercomplex (Fig. 4C) in which a bicylindrical *frl*-PBS docks symmetrically between two PSI trimers. ApcE2 serves as the apparent interaction site, positioned close to the FRL-specific PSI subunit PsaB2 and near PsaD, which is expressed from the WL gene (Fig. 4D). Owing to the limited number of particles, our STA map of the supercomplex does not resolve secondary structural features, and it remains unclear whether additional factors mediate or stabilize PBS-PSI trimer interaction (Fig. 4C). Notably, applying the same strategy to the WL data did not yield solitary PBS, nor were such complexes discernible in manual inspection of WL tomograms. Furthermore, because only *frl*-PSI can form trimers in *C. thermalis*^16,36^, the structure we resolved of a single *frl*-PBS bound to two trimers of *frl*-PSI is likely specific to FRL growth conditions, when the FRL gene cluster is expressed.

**Figure 4.**
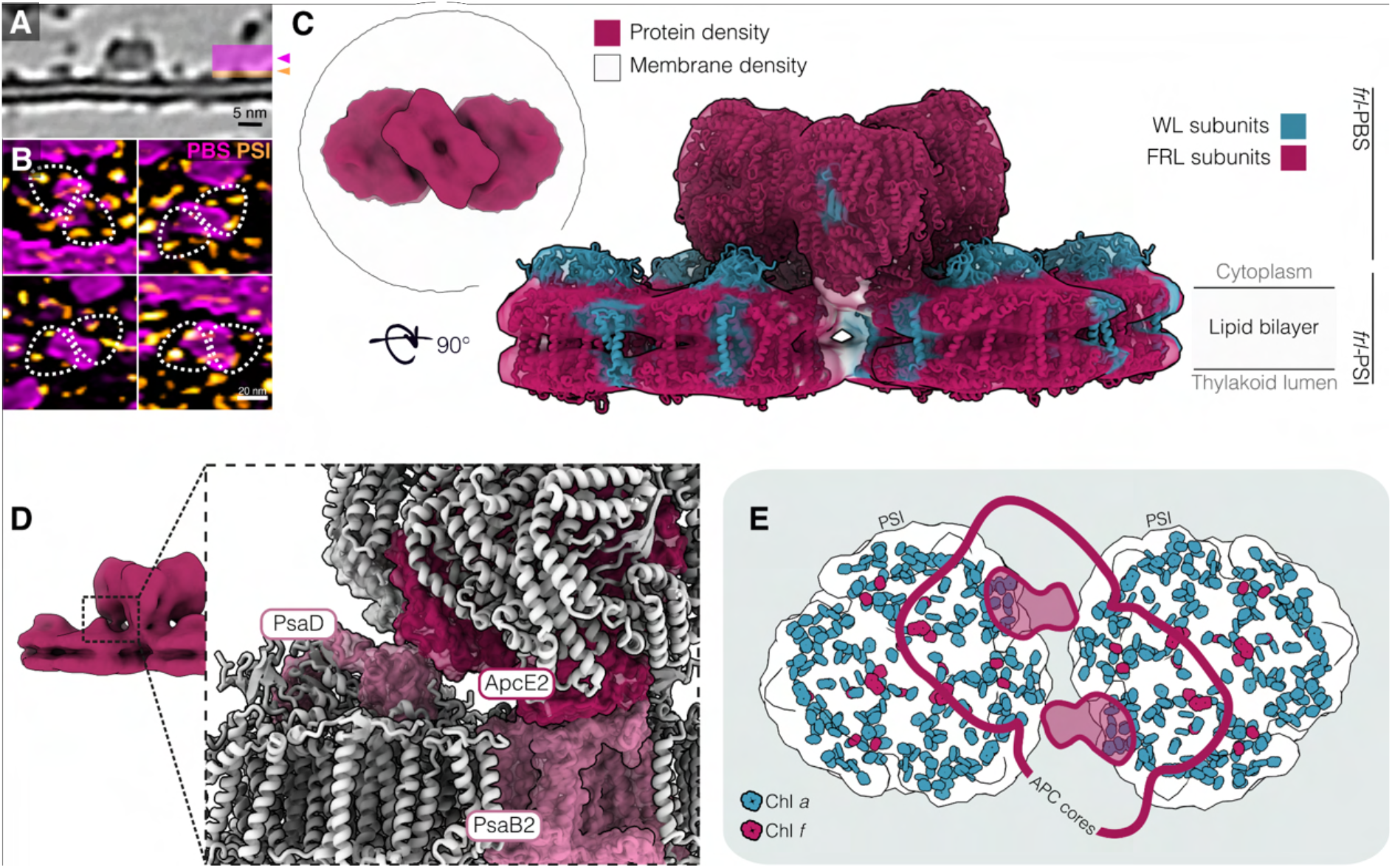
A *frl*-PBS-PSI supercomplex in FRL-acclimated cells. **A**, Tomographic slice showing a solitary *frl*-PBS on the cytosolic surface of the thylakoid membrane, associated with small cytosolic protruding densities. **B**, Surface view overlay of membrane-proximal densities from the same region. Densities at ∼6 nm from the thylakoid surface, corresponding to *frl*-PBS, are shown in magenta, while densities at ∼2 nm from the membrane surface, corresponding to PSI, are shown in orange. Dashed outlines indicate trimeric PSI densities associated with solitary *frl-*PBS. **C**, Subtomogram average of the *frl*-PBS-PSI supercomplex (global resolution: 22.2 Å, **Fig. S13E**). The top-left inset shows a top-view of the complex, with membrane density shown as an outline. The side view shows STA near fitted models of the *frl-*PBS (PDB: 9I1R) and *frl-*PSI (PDB 9EYS). FaRLiP-encoded subunits are shown in red and WL subunits in blue. **D**, Zoom in on the interaction site between PBS and PSI, as seen from the symmetry interface. The PBS interacts though ApcE2, primarily with PsaB2, while also making close contacts with PsaD. The PBS interacting subunit is shown in darker red, PSI interacting subunits are shown in pink. **E**, Schematic top view of the *frl*-PBS-PSI supercomplex. The position of the PBS-APC cores is indicated by the red outline and contact points at the membrane surface are highlighted. Aromatic rings of PSI chlorophylls are shown, with Chl *a* in blue and Chl *f* in red. The docking geometry places terminal bilins at substantial distance from the nearest long-wavelength chlorophyll site.

This *frl*-PBS-PSI organization is distinct from previously described CpcL- or CpcG2-type PBS systems^37,38^, in which PBS connection to PSI is mediated by rod-associated PBS components. Instead, our FRL supercomplex consists of a bicylindrical PBS bridging two *frl*-PSI trimers to form the first described structure of a full PBS in complex with PSI. Interestingly, Chl *f* is not evenly distributed in *frl-*PSI, with most of the red-shifted pigments being confined to one side of the monomer^39^. The *frl-*PBS-PSI supercomplex conformation positions the PBS on the PSI side lacking Chl *f,* placing the closest bilin (K801) at distance of 41.7 Å to the nearest long wavelength absorbing Chl *f* site A21 (Fig. 4E; Fig. S12B). This relatively long distance would be expected to significantly slow down excitation energy transfer, suggesting that without additional pigment-containing proteins, turnover of this supercomplex would be comparatively inefficient.

The low abundance of this *frl*-PBS-PSI supercomplex argues against direct energy transfer from PBS to PSI being the dominant excitation path to PSI under our conditions; many PSI are positioned close to PBS-PSII rows, implying indirect excitation transfer via PSII. However, the presence of this supercomplex inside FRL-adapted cells means that PBS-to-PSI energy transfer can occur through a discrete arrangement where two trimers of PSI share a common antenna, potentially serving a regulatory role.

### *wl*-PBS are retained in FRL

Interestingly, FRL-adapted cells often retained a small reserve of *wl-*PBS (∼7% of all PBS in tomograms) (Fig. S14G). Even cultures grown exclusively in FRL for ∼6 months still consistently maintained some *wl*-PBS (Fig. 5). In order to control for possible outlier effects due to undersampling in cryo-ET, we exploited the characteristic red-shift of chlorophyll autofluorescence in FRL to verify that individual imaged cells were fully acclimated to FRL conditions. We developed a hyperspectral correlative light and electron microscopy (hsCLEM) workflow using a cryo-fluorescence microscope with a white-light laser and prism-based spectral detectors to obtain spatially resolved low-temperature chlorophyll emission spectra that report on the acclimation state of individual cells on an EM grid, which are subsequently imaged by cryo-ET (see Methods; Fig. S15). Using our hsCLEM pipeline, we confirmed that *wl*-PBS are present in cells that show a single red-shifted chlorophyll emission profile peaking at ∼755 nm, as expected for photosystem terminal emitters that are red-shifted^40^, indicating that essentially all photosystems have incorporated Chl *f* and *d* (Fig. 5A-F; Fig. S14). Furthermore, hsCLEM spectra of PBS excitation consistently showed red-shifted emission at 745nm (Fig. S14D,E). STA of these *wl*-PBS in FRL-adapted cells confirmed the structure is indistinguishable from that of *wl*-PBS in WL, and that these *wl*-PBS dock onto PSII in the same asymmetrical manner and dimer-to-dimer spacing as seen in WL (Fig. 5G; S16F). While particle number limitations prevented us from resolving the associated PSII in sufficient detail to unambiguously identify it as *frl-*PSII, this combined data suggests that retained *wl*-PBS are promiscuous, able to associate to *frl*-PSII and transfer excitation energy to the red-shifted cores. Again, this affirms the idea that PBS binding, rather than the variant of PSII, regulates the spacing and geometry of PSII rows.

**Figure 5.**
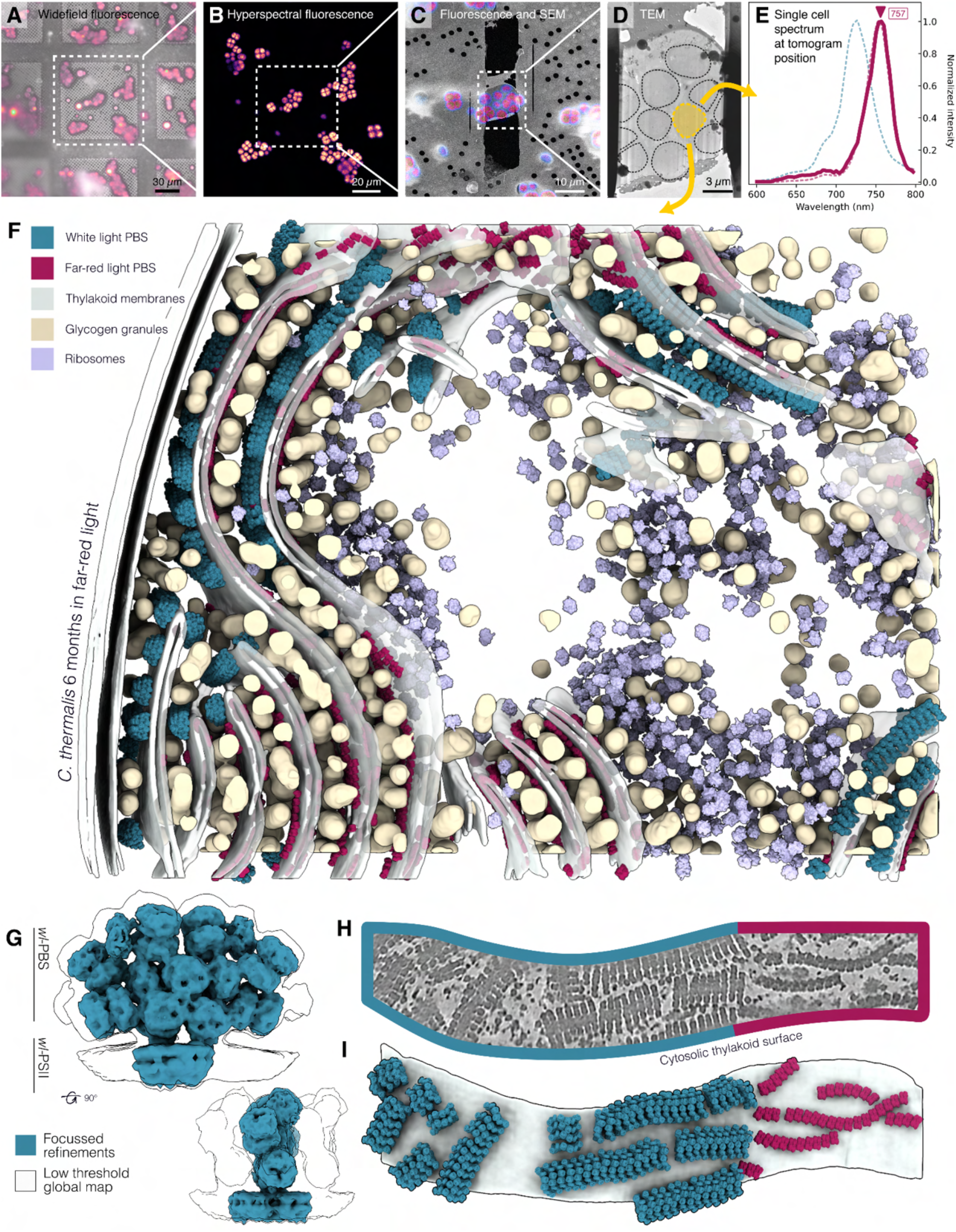
Hyperspectral CLEM reveals that *wl*-PBS persist in fully FRL-acclimated cells. **A–F**, Hyperspectral CLEM workflow links cellular acclimation state to thylakoid ultrastructure in individual, FRL-acclimated *C. thermalis* cells. **A**, Widefield fluorescence image of cyanobacterial cells frozen on an EM grid, with the grid surface shown in gray and chlorophyll autofluorescence (575-615 nm) in magenta. **B**, Hyperspectral fluorescence image of the same cells, acquired as a wavelength scan from 600–800 nm in magenta-to-yellow (Inferno) intensity colormap. **C**, Overlay of the hyperspectral fluorescence image with an SEM image of the corresponding FIB-milled lamella. **D**, TEM overview of the lamella, with cell outlines annotated and the cell selected for analysis highlighted in yellow. **E**, Single-cell chlorophyll emission spectrum extracted from the selected cell, showing a fully red-shifted FRL phenotype. Mean spectra from WL (n = 2.2k) and FRL cells (n = 12.2k) are shown as dotted reference curves. **F**, Segmentation of the tomogram acquired from the highlighted cell. *wl*-PBS are shown in blue, *frl*-PBS in red, thylakoid membranes in light grey, glycogen granules in tan, and ribosomes in lilac. Despite the fully FRL-acclimated spectral state of the cell, *wl*-PBS are retained on the thylakoid membranes together with *frl*-PBS. **G**, Subtomogram average of the retained *wl*-PBS-PSII supercomplex from FRL-acclimated cells. The complex has the same overall architecture as *wl*-PBS-PSII observed in WL-grown cells. **H, I**, Flattened membrane surface view and corresponding map-backs showing *wl*-PBS and *frl*-PBS on the same thylakoid surface. *wl-*PBS and *frl-*PBS do not form heterotypic rows, but rather segregate into distinct membrane domains.

The remaining *wl*-PBS therefore coexist with *frl*-photosystems in cells that have otherwise adapted to the FRL state, and these *wl*-PBS may even associate with *frl*-PSII. Their retention could reflect slow antenna turnover, incomplete repression of WL genes in our light conditions, or the presence of an active mechanism that maintains a residual *wl*-PBS pool. The latter possibility is compelling in the ecological context of FaRLiP cyanobacteria, which often inhabit dense microbial communities where the light spectrum can fluctuate between visible and far-red regimes, and the ability to rapidly revert from FRL to WL photosynthesis could be advantageous.

Critically, the coexistence of *wl*- and *frl*-PBS does not lead to intermixed assembly; we did not observe heterotypic arrays in which *wl*-PBS and *frl*-PBS share a row of PSII (Fig. 5H,I). This may be a result of the distinct PSII-binding geometries between *wl*-PBS and *frl*-PBS, which could prevent incorporation of the other PBS type into the same row (Fig. 3F). While *wl-*PBS and *frl-*PBS rows can co-occupy the same membrane surface, nearest neighbor analysis showed that *wl-*PBS rows were found adjacent to other *wl-*PBS rows three times more frequently than a random distribution despite being the lower abundance species in FRL cells, implying partial segregation into *wl*-PBS or *frl*-PBS domains (Fig. S14F). One explanation for this is that rows of *wl*-PBS accumulate in flat or convex regions of the thylakoids, while *frl-*PBS can distribute evenly across the thylakoid membranes, leading to a separation between the complexes. In addition, this separation of homotypic domains may reflect the complex process of PSII synthesis and assembly in cyanobacteria, which was shown to be concentrated in specific membrane regions^41,42^. Moreover, concerted timing of supercomplex synthesis and assembly could lead to their local accumulation. For example, transient expression of white light genes could promote local synthesis of *wl*-antenna and photosystem components, leading to clustered WL-like domains. Segregation of *wl*- and *frl*-PBS rows could help maintain coherent antenna-photosystem assemblies when cells contain mixed WL and FRL components, preserving consistent energy transfer in FRL and during the process of acclimation.

Together, our results depict cyanobacterial thylakoid membranes as a highly regulated and dynamic system. In green algae and plants, the light harvesting complexes are known to be critical in maintaining the lateral heterogeneity that is needed for spatial regulation of the photosynthetic machinery^43,44^. Similarly, the PBS emerges as more than simply an accessory antenna to the photosystems, but as a key regulator of molecular architecture and the agent of cyanobacterial thylakoid plasticity under changing light conditions. The modular nature of the PBS makes it a key factor not only in efficient energy transfer at the molecular level, but also in directing the spatial organization of photosynthetic complexes and thylakoid architecture at the membrane scale.

## Data availability

Cryo-ET example tomograms and all generated STA maps are available at the Electron Microscopy Data Bank (EMDB) with the accession codes EMD-100373-100391. All custom scripts used for analysis in this work are publicly available in the dedicated GitHub directory: https://github.com/eSveeDee/vanDorst2026_CT-FaRLiP. Scripts for the analysis of hyperspectral CLEM data are packaged in StellarStats: https://github.com/eSveeDee/StellarStats.

## Acknowledgements

Calculations were performed at the sciCORE (http://scicore.unibas.ch/) scientific computing center at the University of Basel, Switzerland. We thank Miroslav Peterek and Bilal Qureshi for access to microscopes and support at ScopeM, ETH Zurich. We also thank the BioEM Lab at the University of Basel for access to microscopes and support. We extend our gratitude to Giovanni Consoli for discussion and Marco Leong for early access to the *frl*-PSII model.

## Methods

### Culture conditions

*Chroococcidiopsis thermalis* PCC 7203 cells were cultured in 50 mL volume in 150 mL flasks in liquid BG11^1^ medium, shaking at 110 rpm and 27 °C. (Fig. S1A) Cultures grown under white light conditions were exposed to continuous white light (WL; 14 µmol photons m⁻² s⁻¹) while far-red light (FRL) cultures were exposed to continuous 730 nm far-red light (20 µmol photons m⁻² s⁻¹) (Fig. S1B). Light intensity was measured at position of the culture using a MQ-500 quantum meter (Apogee). Cultures were initially transitioned from WL to FRL for at least 14 days, or until the amount of Chl *f* stabilized, as measured by whole cell absorption, before being considered FRL-acclimated (Fig. S1C). FRL cultures were thereafter maintained continuously under FRL.

Cells in the logarithmic growth phase were diluted 1:1 into fresh medium 24 h before experiments. For each experiment, to assure no back acclimation to WL occurred, the state of the cell cultures was tested by measuring the relative amount of Chl *f* (710 nm) compared to Chl *a* (678 nm) by absorption spectroscopy using concentrated (∼100×) cell suspensions in a NanoDrop UV-Vis spectrophotometer (DeNovix DS-11). Cultures were then concentrated to an optical density at 600nm (OD600) of ∼3-5 for plunge freezing by gentle centrifugation at 3,000 × *g* at RT for 3 min.

### Plunge freezing

Cells were plunge-frozen using a LEICA EM GP2 plunge freezer (Leica Microsystems). A 3.5 µL aliquot of concentrated cells was applied to glow-discharged R2/1 carbon-coated 200-mesh copper EM grids (Quantifoil). Excess liquid was blotted from the back side for 2 s, and grids were plunged into liquid ethane cooled to approximately −182 °C. Grids were stored in liquid nitrogen until further processing. For cryo-FIB milling, grids were clipped into copper autogrid supports with a modified cut-out geometry (Thermo Fisher Scientific).

### Hyperspectral correlative light and electron microscopy (hsCLEM)

The hsCLEM workflow was used to assign the spectral acclimation state of individual cells before correlating these cells with cryo-FIB lamellae and tomographic data (Fig. 5; S15). Low-temperature spectral fluorescence imaging was performed on a Leica Stellaris 8 microscope (Leica Microsystems) equipped with a white-light laser, HyD S detectors, and a cryo-stage operated at −193 °C. Grid overviews were acquired in a widefield mode using reflected light and autofluorescence (ex: 541-551, em: 565-605) (Fig. S15A). Fluorescence emission wavelength scans were acquired in the *xyλ* mode in the 600-800 nm range, with a 3 nm step size and a 15 nm detection bandwidth (Fig. S15B). Chlorophyll emission spectra were recorded using 440 nm excitation, while phycobilisome emission spectra were recorded using 560 nm excitation. For both channels, laser power was set to 10%. All imaging operations were performed using the Navigator module of LAS X software (Leica Microsystems). The resulting spatially resolved spectral data were analyzed using custom Python scripts implemented in the StellarStats package (https://github.com/eSveeDee/StellarStats).

Briefly, cells were segmented from maximum-intensity projections of wavelength scans using Cellpose3^2^ (Fig. S15C). Segmented cell masks were used to extract the mean fluorescence intensity of each cell at each wavelength (Fig. S15D). Cells with a raw maximum intensity below 5, or with peak wavelengths outside the expected physiological range (<680 nm for chlorophyll spectra or <615 nm for PBS spectra), were discarded as false-positives. To combine experiments with slight offsets in detection range, spectra sampled at 3 nm intervals were linearly interpolated to 1 nm spacing. Single-cell spectra were normalized to the terminal emitter peak in the 710-780 nm range. For mean spectra, all segmented cells passing the abovementioned quality filters were grouped by condition and averaged after normalization (440 spectra: n=12204 WL, n=2291 WL, 560 spectra: n=1557 FRL, n=2429 WL) (Fig. S14D-E). These WL and FRL reference spectra were used to assess whether individual cells selected for tomography showed a fully FRL-acclimated phenotype. Cells analyzed by cryoET were manually correlated to their corresponding fluorescence spectra by consecutive overlays of confocal fluorescence, widefield fluorescence, SEM, and TEM images of the lamella site.

### Focused ion beam (FIB) milling

Cryo-FIB milling of the samples was performed following the protocol established in Schaffer et al.^3^ using an Aquilos 2.5 dual-beam FIB/SEM (Thermo Fisher Scientific). The scanning electron beam was used to acquire grid overviews that were overlayed with the widefield grid maps collected at the fluorescence microscope. Where applicable, milling was performed at sites of spectral LM acquisition. Grids were coated with a layer of organometallic platinum ahead of milling using the gas injection system (GIS). The samples were milled with a gallium ion beam, using progressively smaller milling patterns and milling currents until a thickness of 100-200 nm for the final lamellae was reached. Grids were then stored in liquid nitrogen until acquisition at the TEM.

### CryoET data acquisition

Tomographic data were acquired on a 300 kV Titan Krios G3 microscope (Thermo Fisher Scientific) equipped with a post-column BioQuantum energy filter operated with a 20 eV slit width and a K3 direct electron detector (Gatan). Tilt series were acquired using SerialEM^4^ software with a dose-symmetric tilt scheme starting at the lamella pre-tilt angle (−10°), with 2° increments over a total angular range of 112-116°. Tilt projections were recorded in counting mode into 7-13 frames, with a total dose of approximately 120 e⁻ Å⁻² per tilt series and a calibrated pixel size of 2.6 Å. Target defocus values ranged from 2.5-5 µm. Additional WL data was acquired on a 300 kV Titan Krios G4 microscope equipped with a Falcon 4i direct electron detector and a Selectris X energy filter (Thermo Fisher Scientific) operated with a 10 eV slit width. Tilt series were acquired using Tomography software (Thermo Fisher Scientific) with a dose-symmetric tilt scheme starting at the lamella pre-tilt angle (−10°), 2° increments, and a total angular range of 112°. Tilt projections were recorded as EER movies with 270 frames per movie, a total dose of approximately 120 e⁻ Å⁻², and a calibrated pixel size of 2.34 Å. Target defocus values were in the 2.5-5 µm range.

### Tomogram preprocessing, reconstruction and denoising

Tomogram preprocessing and reconstruction followed the general workflows for cryoET reconstruction using Scipion^5,6^ environment and AreTomo3^7^ described in TomoGuide^8^. For K3 data, preprocessing and reconstruction were performed within the Scipion framework. Raw tilt series were motion-corrected using MotionCor2^9^, bad tilts were manually excluded, and tilt-series alignment was performed using patch tracking in Etomo/IMOD^10,11^. CTF estimation was performed using CTFFIND5^12^, and 2D CTF correction was performed using IMOD. For Falcon 4i data, preprocessing, motion correction, CTF estimation, tilt-series alignment, and reconstruction of CTF-corrected tomograms were performed using AreTomo3^7^ (Fig. S4).

Tomograms were denoised and missing-wedge-corrected using DeepDeWedge^13^ on tomogram pairs reconstructed from odd and even frames.

### Data visualization

Membranes were segmented in denoised tomograms using MemBrain-seg^14^ and used as membrane masks to create flattened membrane renderings using MPicker^15^. Membrane surfaces were selected using a single seed point and flattened. To generate comparative renderings of protein complexes at defined distances from the membrane surface (Fig 2, 4A; S5; S12A), 2-3 slices were summed at selected offsets relative to the membrane surface (on the cytosolic side: 6 nm for PBS, 2nm for PSI, on the luminal side: 2nm for PSII). These projections were overlaid in false color using Fiji (v2.16)^16^. PSI and PSII densities on the cytosolic side of the thylakoid membrane were assigned based on their size, position near the membrane, and characteristic organization.

EM maps and segmentations were visualized in ChimeraX^17^. For 3D renderings of complete tomograms (Fig. 1; S2), membranes were segmented using MemBrain-seg and manually corrected in Amira3D^18^. Glycogen granules were segmented using a custom-trained model in Ais^19^. Particles identified by template matching and subtomogram averaging were mapped back into tomographic reconstructions, and all components were visualized together using the ArtiaX^20^ plugin in ChimeraX.

### Subtomogram averaging

Subtomogram averaging workflows are summarized in Fig. S7. To generate initial maps for *wl*-PBS and *frl*-PBS particle picking, MemBrain-seg and MPicker were used to create flattened thylakoid membrane surfaces as described above. Points were manually picked along PBS rows in MPicker for three tomograms, ensuring that the particle normal angle was assigned toward the cytoplasmic side and that the second angular direction followed the row direction. These positions were used to fit splines along picked rows, and particles were interpolated every 64 Å using STOPGAP^21^ (Fig. S7A). Angle priors were then reassigned to the interpolated particles using a custom Python script, with one axis aligned to the membrane normal and the in-plane angle aligned to the median row direction of the manually assigned coordinates. Subtomograms were extracted at these positions at bin4 (10.4 Å/px) using STOPGAP (Fig. S7B). Particles were aligned with local searches to approximately 30 Å resolution. These maps were then used as templates for 3D template matching with pytom-match-pick^22^ (Fig. S7C). Template matching was performed on bin4, CTF corrected tomograms, with 10° angular sampling. The particle extraction threshold was manually adjusted for each tomogram to minimize false positives. Manually generated boundary masks and MemBrain-seg membrane segmentations were used to exclude membranes and vacuum from the search volume. Picks were imported into RELION-5^23^ and processed using the subtomogram averaging workflow described in TomoGuide^8^. Two rounds of template matching and alignment were performed to generate the final templates.

### *wl-*PBS-PSII

Template matching was performed with pytom-match-pick on bin4 tomograms (9.36 Å px⁻¹) using high angular sampling (4.9°) and twofold symmetry around the z-axis. A data-derived template of wl-PBS in a row was used, with a focused mask on the central PBS (Fig. S7E-F). Lamella boundary masks in combination with pytom-match-pick’s default cut-off estimation were used to extract particle positions.

A custom Python script was used to assign a unique row number (row ID) to each PBS row (Fig. S7G). Briefly, particle coordinates and angles were treated as a point-cloud system, and rows were identified by connected-component analysis of a nearest-neighbor graph. Edges were defined using a spatial threshold of 200 Å and angular difference thresholds for rot, tilt and psi angles (20°, 10°, 20° respectively). To fill gaps in rows, one round of dilation and erosion of particle positions was performed along the row axis. Singleton particles were discarded. A 3D distance cleaning was performed at a 130 Å cut-off. Particle half-sets were assigned by row ID to maintain independent half-sets for gold-standard FSC measurements. This resulted in 171046 particles across 79 tomograms, which were imported into RELION5 and extracted at bin4 (9.36 Å px⁻¹) (Fig. S7H). These particles were subjected to a round of 3D classification, without mask and alignment ( T=0.01, K=5, 63176 particles). Selected particles were aligned in Refine3D with local angular searches (3.7°, --sigma_ang 3) with a mask covering two PBS and two PSII, to a resolution of 19.4 Å. A bin1 reference (2.34 Å px⁻¹) was generated for CTF refinement and Bayesian polishing, improving the overall resolution to 9.3 Å. Particles were re-extracted at bin2 (4.68 Å px⁻¹) and subjected to another round of 3D refinement, yielding a map at 9.2 Å. A further round of 3D classification (T=1, K=4;), masked for the membrane domains. Two opposing orientations of PSII were found (20305 particles in one orientation and 22067 in the other), as dictated by the asymmetry of the conventional cyanobacterial PBS-PSII complex. The particles were rotated to a shared orientation and subjected to another round of Refine3D with local searches, followed by CTF refinement and Bayesian polishing yielded a map clearly resolving PBS and PSII at 9.4 Å. Re-extraction at bin1 and refinement resulted in a final 9.1 Å global resolution map of the two-PBS two-PSII masked supercomplex. At this stage, a local-resolution-filtered map of the full supercomplex was generated to represent the overall organization of the row-associated *wl*-PBS-PSII assembly.

For focused refinement of *wl-*PBS, the central PBS was masked for an additional round of Refine3D, improving the map resolution to 8.0 Å. Finally, C2-symmetry was applied in the last round of alignments, resulting in a final *wl-*PBS map with a resolution of 7.2 Å.

*wl-*PBS was further aligned from the previously aligned supercomplex with initially a mask focused on two central PSII dimers. These were subjected to a round of classification (T=10, K=3, 29773 particles) to select for full PSII occupancy and remove row-ends and remaining false positives from the dataset. The particles were then centered on the most central PSII dimer in the xy plane to allow imposing C2-symmetry in the succeeding Refine3D alignment, resulting in a 11.5Å map. A final round of classification was applied to select for the best particles (T=20, K=3, 14315 particles), which after a final alignment with C2-symmetry resulted in a 9.9Å resolution map, as reported by gold-standard FSC (Fig. S6K).

For the averaging of row-ends, the bin1-aligned supercomplex reconstruction was re-extracted in bin4, and subjected to two rounds of classification to select a class of row-ends (T=1, final 8465 particles) (Fig. S11A). In the row end class, an additional subunit appeared on the phycobilisome in the Rs1* position (Fig. S11D). A last round of classification with a focused mask (Fig. S11C) was applied to differentiate two distinct classes: one row-end with the trailing Rs1 in the regular position (Fig. S11B) representing 45.5% of the particles, and a class with repositioning of the Rs1 toroid into the Rs1* position (Fig. S11D), representing 35.7% of particles. The remaining particles (18.8%) present in ambiguous classes, likely represent a continuous conformational space between the two major populations.

### *frl*-PBS-PSII

Template matching was performed on bin4 tomograms (10.74 Å px⁻¹) using high angular sampling (4.9°) and two-fold symmetry around the z-axis. A data-derived template of *frl*-PBS in a row was used, with a focused mask on the central three PBS to enhance the cross-correlation signal (Fig. S6E-F). Manually generated boundary masks and MemBrain-seg membrane segmentations were used to exclude membrane and vacuum from the search volume. pytom-match-pick’s automatic extraction threshold was used to extract particle positions.

The nearest-neighbor graph workflow described above was used to assign row IDs, fill gaps in rows, remove singletons and duplicates, and assign row-based half-sets, resulting in 125,294 particles from 29 tomograms (Fig. S6G). Particles were imported into RELION5, extracted at bin8 (20.8 Å px⁻¹), and subjected to one round of maskless 3D classification without alignment (T=0.5, K=4; 61265 particles) (Fig. S6I). Selected particles were aligned using Refine3D with local angular searches (3.7°, 3.7°, 2 px), extracted at bin4 (10.4 Å px⁻¹), and subjected to further refinement and classification using a mask focused on the central three PBS (T=0.5, K=4; 24,733 particles). After false positives were removed by classification, another round of Refine3D was performed, yielding a 22.18 Å map.

A bin1 reference (2.6 Å px⁻¹) was generated for CTF refinement and Bayesian polishing, improving the overall resolution to 14.1 Å. Particles were re-extracted at bin2 (5.2 Å px⁻¹) and subjected to another round of 3D refinement (1.8°, 1.8°, 3 px), yielding a map at 11.3 Å. A further round of 3D classification (T=0.01, K=5; 23824 particles), masked on the central three PBS, followed by CTF refinement and Bayesian polishing yielded a map at 11.2 Å. Re-extraction at bin1 and refinement (1.8°, 1.8°, 3 px) resulted in a 10.6 Å map of the three-PBS masked supercomplex. At this stage, a local-resolution-filtered map of the full supercomplex was generated to represent the overall organization of the row-associated *frl*-PBS-PSII assembly.

For focused refinement of *frl*-PBS, a mask around the central PBS was used for an additional round of 3D refinement, improving the map to 9.4 Å. A final classification without alignment focused on the central PBS (T = 0.01, K = 5; 13016 particles) was used to select high-quality particles. Subsequent refinement with C2 symmetry, CTF refinement, Bayesian polishing, and refinement using BLUSH regularization yielded a final *frl*-PBS map at 8.2 Å (Fig. S6L).

For focused refinement of *frl*-PSII, particles were refined using a mask on the central two PSII dimers (1.8°, 1.8°, 3 px), yielding an 11.5 Å map. An additional classification (T=0.01, K=5) resulted in a final subset of 2343 particles. These particles were first refined with a mask on two PSII dimers and then with a mask on a single PSII dimer (1.8°, 1.8°, 3 px). Particles were recentered on the selected PSII dimer and re-extracted at bin1. A further refinement was performed with C2 symmetry, using the --sigma_ang flag to restrict alignment to local searches, yielding an 11.2 Å map. A final classification (T=1, K=5, with 3.7° local searches) was used to select the highest-quality particles, and a final round of refinement (3.7°, 3.7°, 2 px, sigma_ang=2) yielded an 8.9 Å map (Fig. S6M).

### *frl*-PBS-PSI

Template matching was performed on the best 7 tomograms using the strategy described for *frl*-PBS-PSII rows. The same data-derived *frl*-PBS-PSII row template was used, but the mask was focused only on the central PBS to ensure strong cross-correlation with solitary *frl*-PBS (Fig. S13A-B). The nearest-neighbor graph workflow was again used to assign row IDs and fill gaps in rows. Remaining singleton PBS particles were now selected by isolating rows with length equal to 1, resulting in 9,665 particles (Fig. S13C). The initial reconstruction yielded an average resembling *frl*-PBS-PSII in a row.

Particles were extracted in RELION5 at bin4 (10.4 Å px⁻¹), and two rounds of maskless classification without alignment (T=0.01, K=5) were used to separate solitary PBS from PBS in rows (Fig. S13D). This yielded a subset of 1204 particles. Reconstruction of these particles revealed a PBS associated with two PSI trimers. Refinement was performed using a mask around the full supercomplex (3.7°, sigma_ang=3), after which the aligned particles were cleaned up with another two rounds of classification (T=0.01, 597 particles final). A bin1 reference was generated for CTF refinement and Bayesian polishing, and particles were re-extracted in bin3 (7.8 Å px⁻¹). Because bin3 alignments did not improve the resolution, another round of CTF refinement and Bayesian polishing as well as a final bin1 refinement were performed, yielding a final map at 22.2 Å (Fig. S13E).

*wl*-PBS in FRL cells

As in *wl-*PBS, the final *wl-*PBS map was used as a template with a mask focused on the central PBS, excluding PSII as not to bias the search (Fig. S16A-B). A similar processing pipeline was applied for *wl-*PBS, using the nearest-neighbor graph network to extract *wl-*PBS rows, resulting in a particle list of 667 particles across 29 FRL tomograms (Fig. S16C). Particles were then extracted in RELION5 in bin4 (10.4 Å px⁻¹), and false positives were removed by a round of classification without mask and without alignment (T=0.1, K=5, 3276 particles). After this, particles were subjected to local refinements in Refine3D, another round of classifications and refinements with a mask focused on three central PBS, resulting in a final particle count of 2737. A round of CTF refinement and Bayesian polishing were applied, after which the particles were reconstructed and aligned with local searches in bin2 (5.2 Å px⁻¹). At this point, a 3D classification with a mask focusing on the membrane was used to separate two opposing PSII orientations (T=1, K=4, 927 and 966 particles in each orientation respectively). Particles were then re-oriented to the PSII orientation and aligned again in bin2. Another round of CTF refinements and Bayesian polishing was followed by a refinement in bin1 (2.6 Å px⁻¹) focused on the central PBS, finalizing the supercomplex average to 18.1 Å resolution as determined by gold-standard FSC (Fig. S16D). To compare PBS-PSII organization of the *wl-*PBS in FRL with WL and FRL supercomplex structures, rigid body fitting of 20 Å maps simulated from atoms was performed in ChimeraX on a local-resolution filtered map. Only A and A’ APC cylinders were included to prevent score bias by PBS size. Correlation of mean data value with the *frl*-PBS-PSII was, 0.5063, while correlation with the *wl*-PBS-PSII was 0.5518, quantitatively confirming that the complex is in the WL confirmation (Fig. S16E-F).

### Spatial analysis

The number of thylakoids from the cell periphery as reported in (Fig. S3A) was manually counted all tomograms with a clear cell-periphery within the field-of-view. Thylakoid stacks were considered up to the central cytoplasmic space. In the case of fascicular architecture, thylakoid up to the center of the spherical formations was considered. Inter thylakoid distances were calculated using Morphometrics^24^. Briefly, Membrain-seg membrane segmentations were manually cleaned and separated using Amira3D and converted into an isotropic mesh. Distances from each vortex to the next thylakoid membrane were calculated. Only inter-thylakoid distances were considered, and high-curvature regions were excluded. Per-membrane median values were taken as single data points and compared between WL and FRL conditions (Fig. S3B).

A point-cloud based spatial analysis was the basis for spatial analysis of PBS organization. Template matching-derived particle lists after cleaning (initial 3D classification, ref step in workflow fig) were subjected to the nearest-neighbor graph workflow described in *Subtomogram Averaging* (Fig. S6G), and individual rows were identified. Radial distance distributions were calculated for each particle to the members of its row (Fig. S3C) resulting in a mean center-to-center distance to the first neighbor of 14 nm for *wl-*PBS and *frl-*PBS. Relative angular deviation of particles in comparison their first row-neighbor was calculated from this and plotted by Euler angle (Fig. S3E). To calculate inter-row spacing, row centroids were calculated for each row as the mean coordinate of each particle, and nearest neighbors were calculated for each row.

To analyze the organization and abundance of *wl*-PBS in FRL conditions only *wl*-PBS and *frl*-PBS particles positions cleaned by classification were compared. Lists were screened for duplicates. Positions within 60 Å in both lists were assumed *wl-*PBS since the *wl-*PBS template did not score against *frl-*PBS. For each tomogram, particles were grouped into rows as described previously, and a 3D nearest neighbor network (k=80) was retrieved (Fig. S14F). The top 3 nearest neighbors within 2500 Å in-plane (within 30° psi-angle deviation) with the particle of origin were considered. The nearest neighbor network was tested for robustness by permutation of search radius, neighbor depth, and angular restriction. For every *wl-*PBS particle, the number of neighbors sharing the same type (*frl* or *wl*) were counted. This value was compared to the same statistic after random permutation of row identity (*frl* or *wl* identity) where *frl* to *wl* ratios were maintained. This was repeated 5000 times to derive a randomized row distribution statistic, which could be compared with the experimental value, resulting in an observed ∼3x enrichment of *wl-*to-*wl* row neighbors.

### Supercomplex model fitting and analysis

To compare supercomplex geometry and pigment distances, composite models were assembled by rigid-body fitting of molecular models into respective subtomogram averages. Focused refinements of PBS, PSII, and PSI were fitted into their respective positions in local-resolution-filtered maps. For *wl*-PBS-PSII, the *Anabaena* sp. PCC 7120 PBS structure (PDB: 7EYD^25^) and the *Synechocystis* sp. 6803 substr. Kazusa PSII structure (PDB: 7N8O^26^) were fitted. For *frl*-PBS-PSII, C. thermalis *frl*-PBS (PDB: 9I1R^27^) and *frl*-PSII (PDB: 9T5T^28^) structures were fitted. For *frl*-PBS-PSI, the C. thermalis *frl*-PSI structure (PDB: 9EYS^29^) was fitted.

Using these composite models, edge-to-edge distances were calculated between phycocyanobilins, defined as donor pigments (CYC), and chlorophylls, defined as acceptor pigments (CLA for Chl *a* and F6C for Chl *f*) (Fig. S10; S16B). Supercomplex geometries (Fig 3F) were determined by generating 20 Å-resolution molmaps of the two lower APC cores and associated PSII or PSI complexes, followed by calculation of their positions and angles from molecular outlines and centers of mass.

Pairwise sequence alignments of far-red light ApcE2 (UniProt K9TUP3) and white light ApcE (UniProt K9U4G6) were calculated using Clustal Omega^30^, via UniProt’s built-in alignment tool with default parameters^31^. The Clustal Omega percentage identity matrix was used to display identity over sequence domains (Fig. S9).

## Supplementary figures

**Figure S1.**
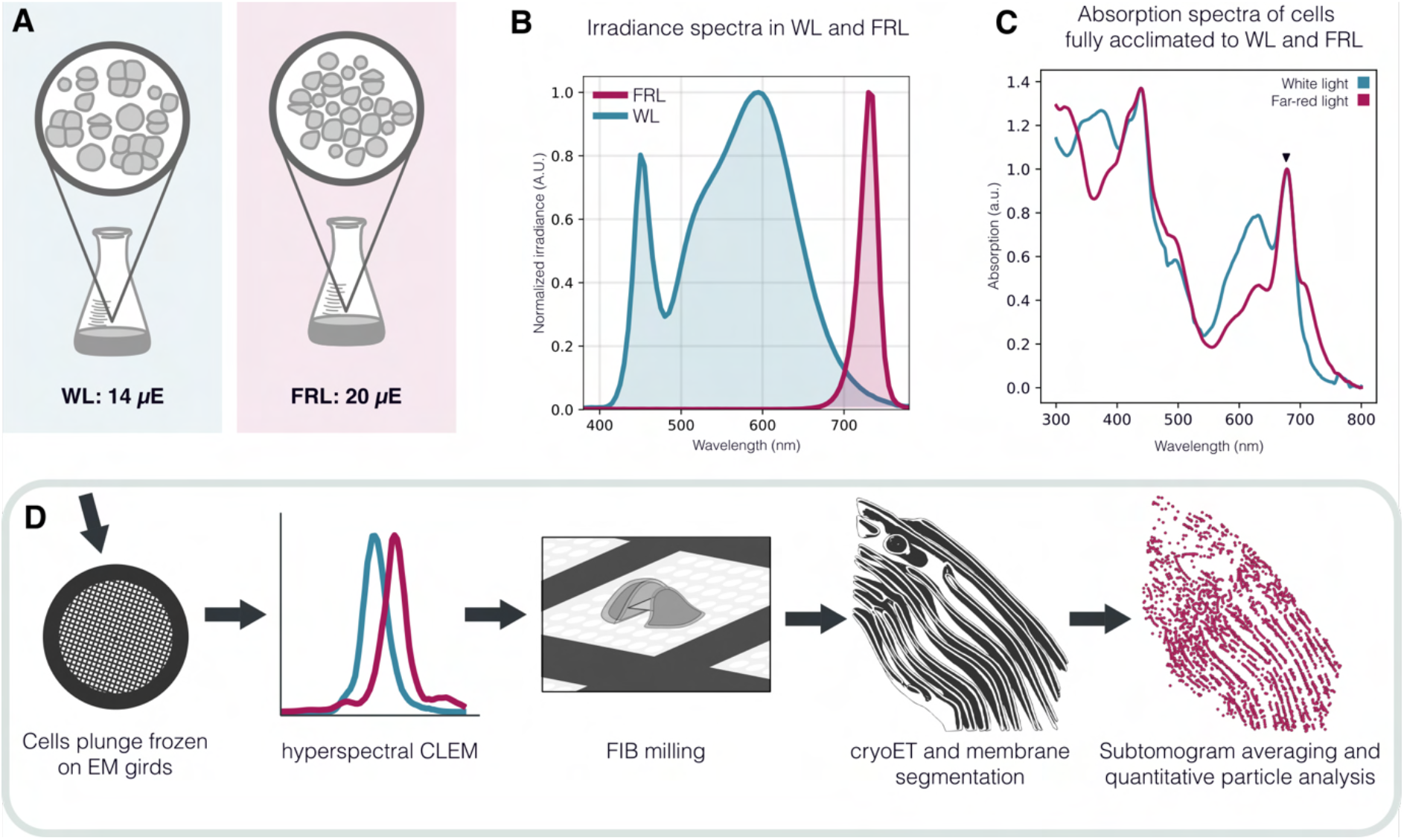
Cell culture and sample preparation overview. **A**, For all the experiments, *C.thermalis* cells were grown in liquid cultures in either White Light (WL, intensity=14uE) or Far-Red Light (FRL, intensity=20uE). **B**, Irradiance spectra of WL and FRL light sources used for culturing. **C**, Representative absorption spectra of concentrated cell culture, measured at room temperature, of cells grown in WL and FRL conditions. **D**, Schematic representation of our cryo-ET workflow. Cells from respective cultures were plunge frozen on EM grids, their fluorescence spectra recorded using hsCLEM, and finally processed in the cryo-ET pipeline to produce tomographic volumes, segmentations, and subtomogram averages (details in Methods).

**Figure S2.**
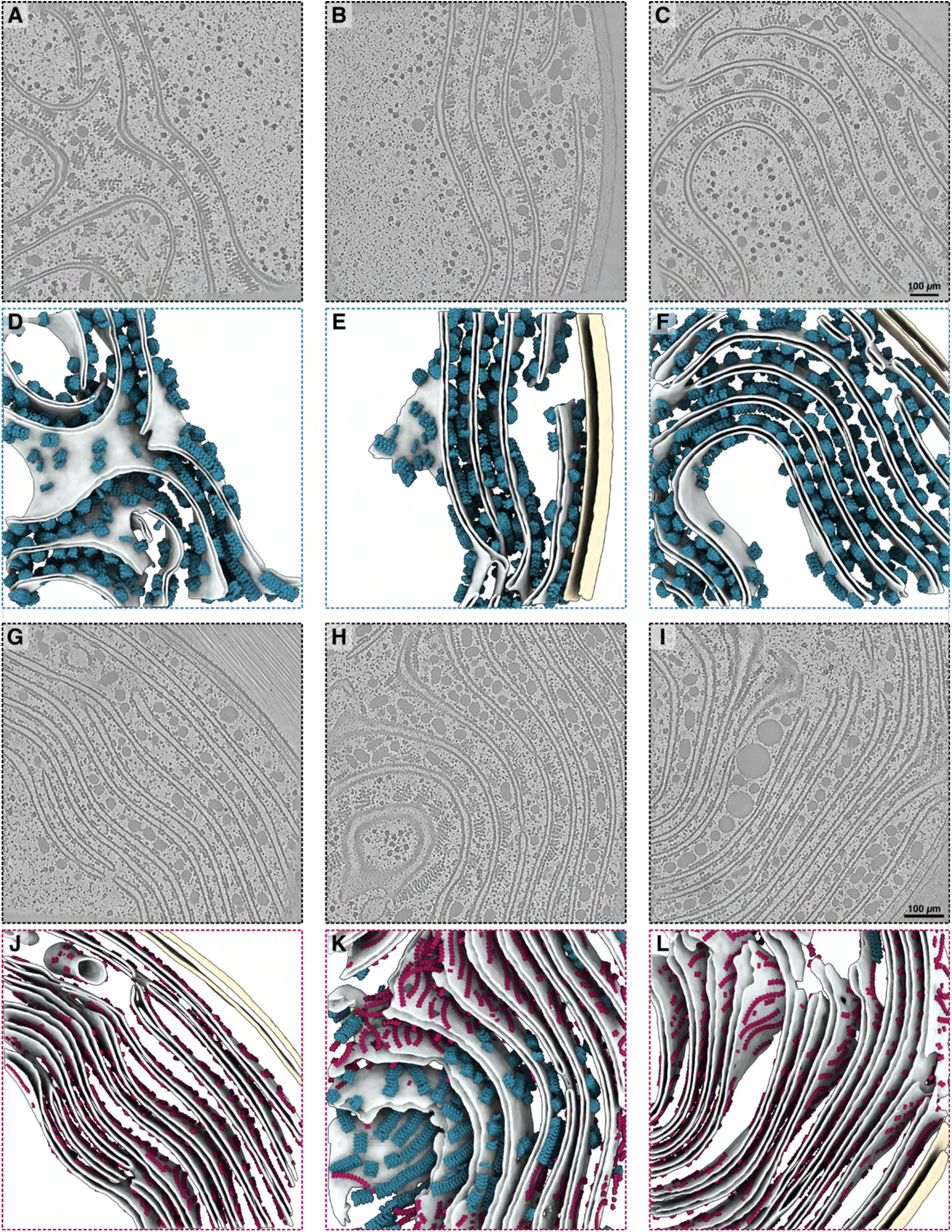
Gallery of tomograms and segmentations from WL and FRL. **A, B, C**, Slices from three tomographic volumes of WL-acclimated *C. thermalis* cells. Scale bar = 100 nm applies to all three panels. **D, E, F**, Corresponding tomogram segmentations and particle map-ins. *wl*-PBS in blue, thylakoid membranes in light gray, cell membranes in light yellow. **G, H, I**, Slices from three tomographic volumes of FRL-acclimated *C. thermalis* cells. Note the higher number of thylakoids compared to WL. Scale bar = 100 nm applies to all three panels. **J, K, L**, Corresponding tomogram segmentations. In all segmentation panels: *frl*-PBS in red, *wl*-PBS in blue, thylakoid membranes in light gray, cell membranes in light yellow.

**Figure S3.**
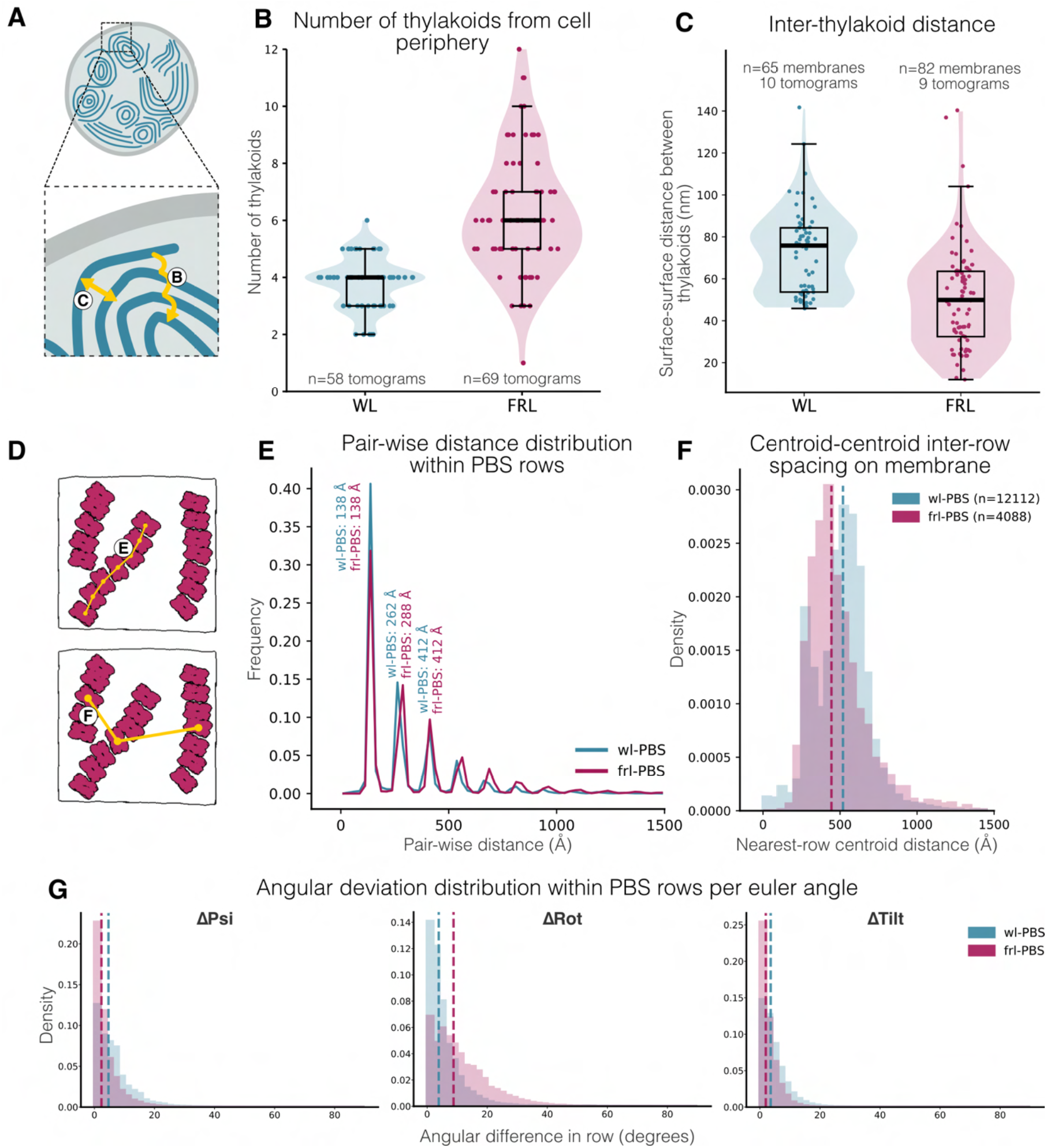
Ultrastructural statistics of thylakoids and PBS rows in WL and FRL. **A**, Schematic representation of thylakoid architecture of C.thermalis and logic for basic thylakoid measurements plots shown in B and C. **B**, Number of thylakoids counted from the cell periphery in WL (mean: 4.3) and FRL (mean: 6.2). **C**, Distance between individual thylakoids in WL (mean: 69 nm) and FRL (mean: 47 nm) measured between surfaces of neighboring membranes. **D**, Schematic representation of PBS rows on the thylakoid membrane as basis for the distance measurements shown in E and F. **E**, Plot of the pair-wise distances between individual PBS within rows in WL and FRL, showing no apparent difference in PBS spacing. **F**, Histogram of distances between PBS rows in WL and FRL measured between centroids of neighboring PBS. Dashed lines show the median values, WL = 52 nm, FRL = 44nm. **G**, Angular distribution of PBS rows in three Euler angles in WL and FRL. Dashed lines show mean values WL = 8.7, 4.6, 7.4 (psi,rot,tilt) and FRL = 3.7, 2.8, 10.9 (psi,rot,tilt), showing there is more rotational freedom along the length of the row in FRL.

**Figure S4.**
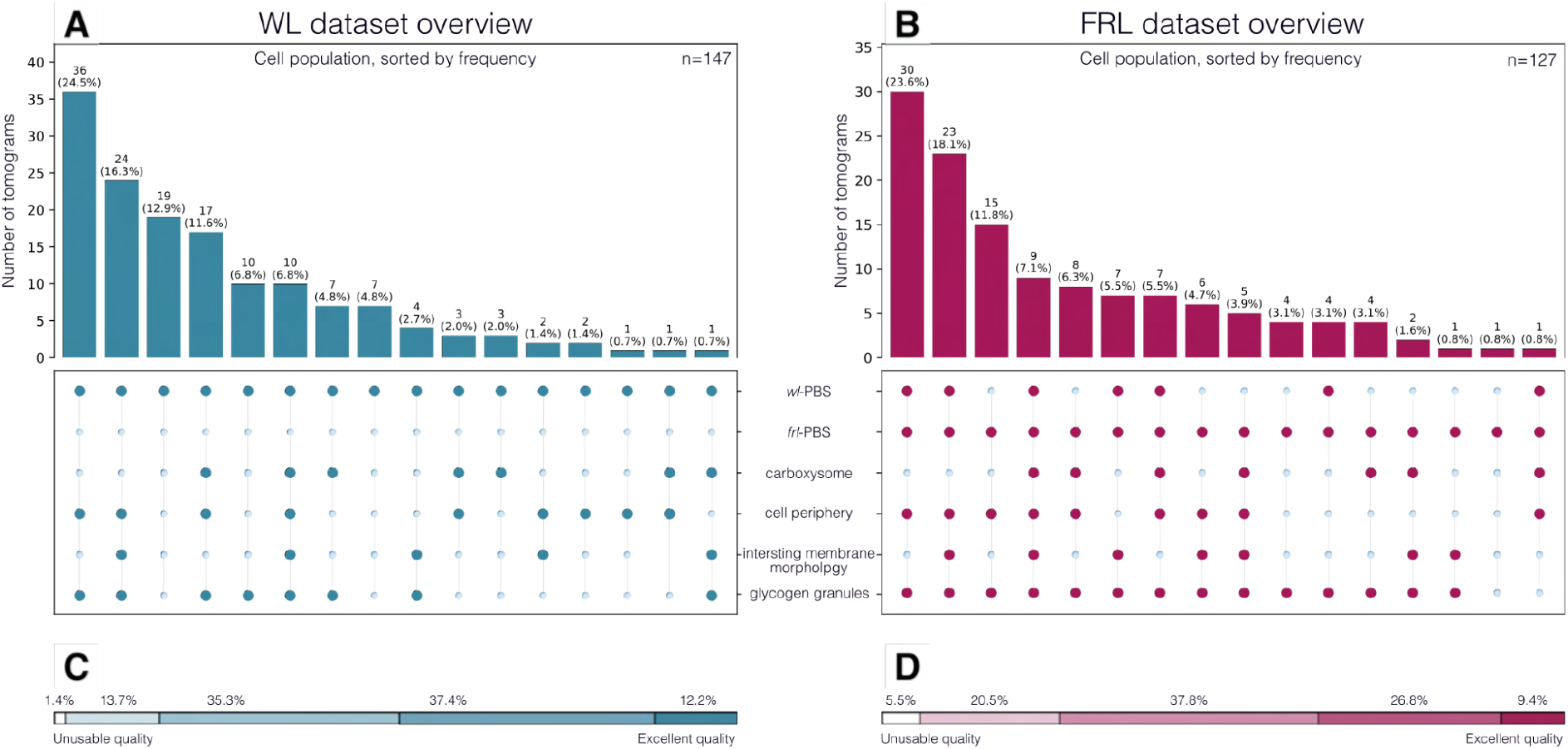
Tomogram dataset statistics. **A, B**, Overview of structural and cellular features present in the complete datasets of WL tomograms n=147 and FRL tomograms n=127. **C, D**, Quality metric assigned for the reconstructed tomograms. Evaluation based on manual inspection of the data.

**Figure S5.**
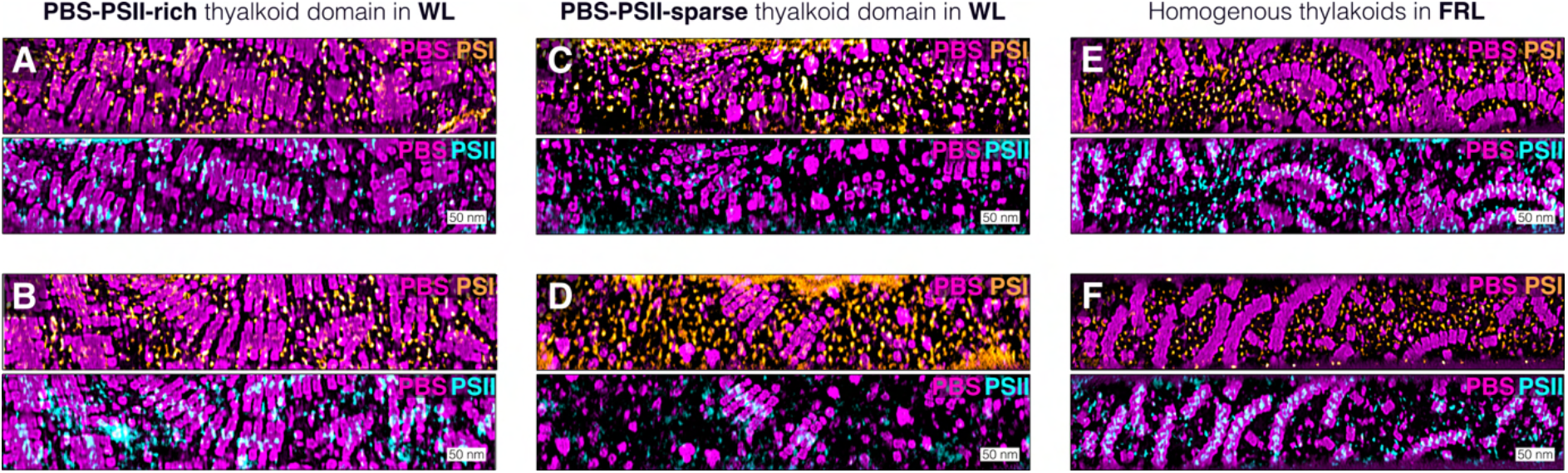
Additional examples of PBS-PSII/PBS-PSI localization on membrane surfaces. **A, B**, Overlays of densities in the PBS-PSII-rich thylakoid regions seen in WL. **C, D**, Overlays of densities in the PBS-PSII-sparse thylakoid regions seen in WL. **E, F**, Overlays of densities in the more homogeneous thylakoids most often seen in FRL. In all panels, PBS are shown in magenta, PSI in orange, and PSII in blue. Scale bars 50 nm.

**Figure S6.**
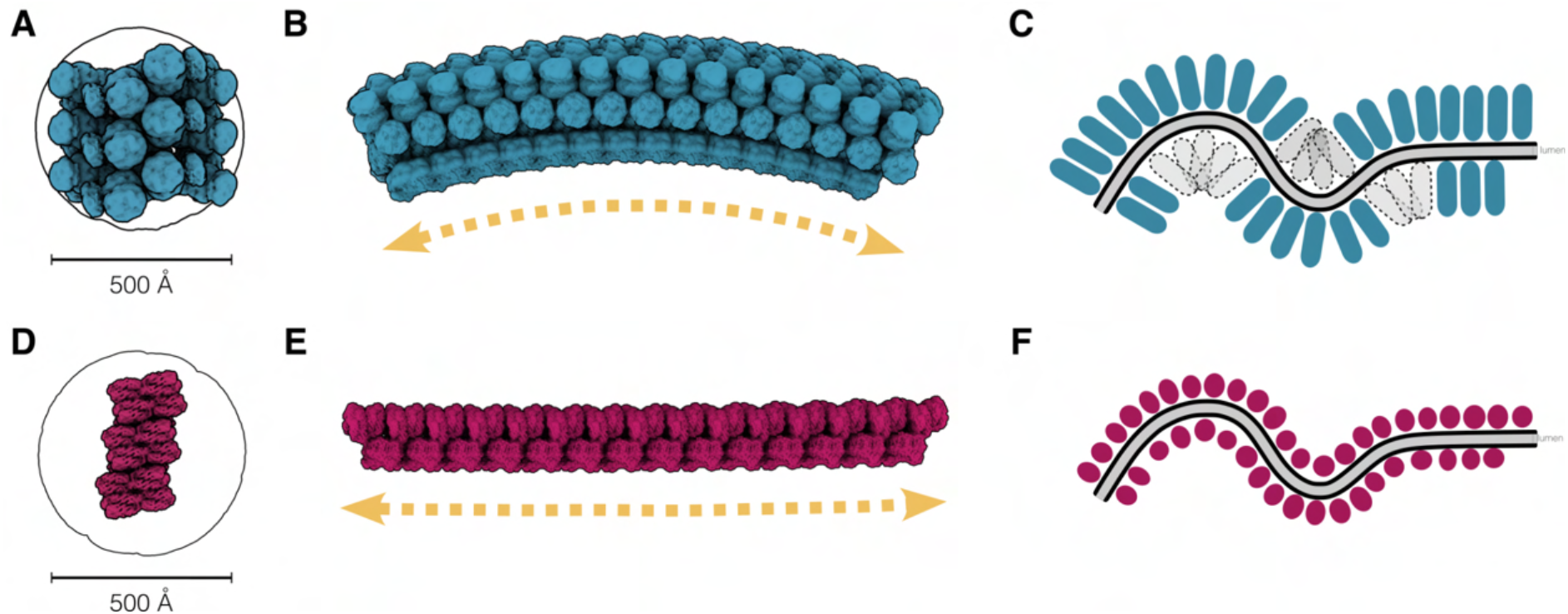
Mesoscale properties of *wl*- and *frl*-PBS. **A**, Top view of a three-PBS containing section of a *wl*-PBS row, demonstrating the membrane footprint of the assembly. **B**, Average row structure, extrapolated from subtomogram average including all *wl*-PBS particles, shows a slightly convex curvature of a *wl*-PBS row. **C**, Overall curvature of the *wl*-PBS rows seen in B can be explained by the localization of the *wl*-PBS on the thylakoid membranes. The large size of the PBS prevents it from occupying concave thylakoid surfaces, leading to an overall convex curvature of the row average. **D**, Top view of a three-PBS piece of a *frl*-PBS row demonstrating the significantly smaller membrane footprint of the FRL assembly compared to WL. **E**, Average row structure, extrapolated from subtomogram average including all the *frl*-PBS particles, is a nearly straight row. **F**, The smaller size of the *frl*-PBS allows it to populate the full physiological curvature range of the thylakoid surface, leading to a homogenous average (a straight row because all the angles average out) as seen in **E**.

**Figure S7.**
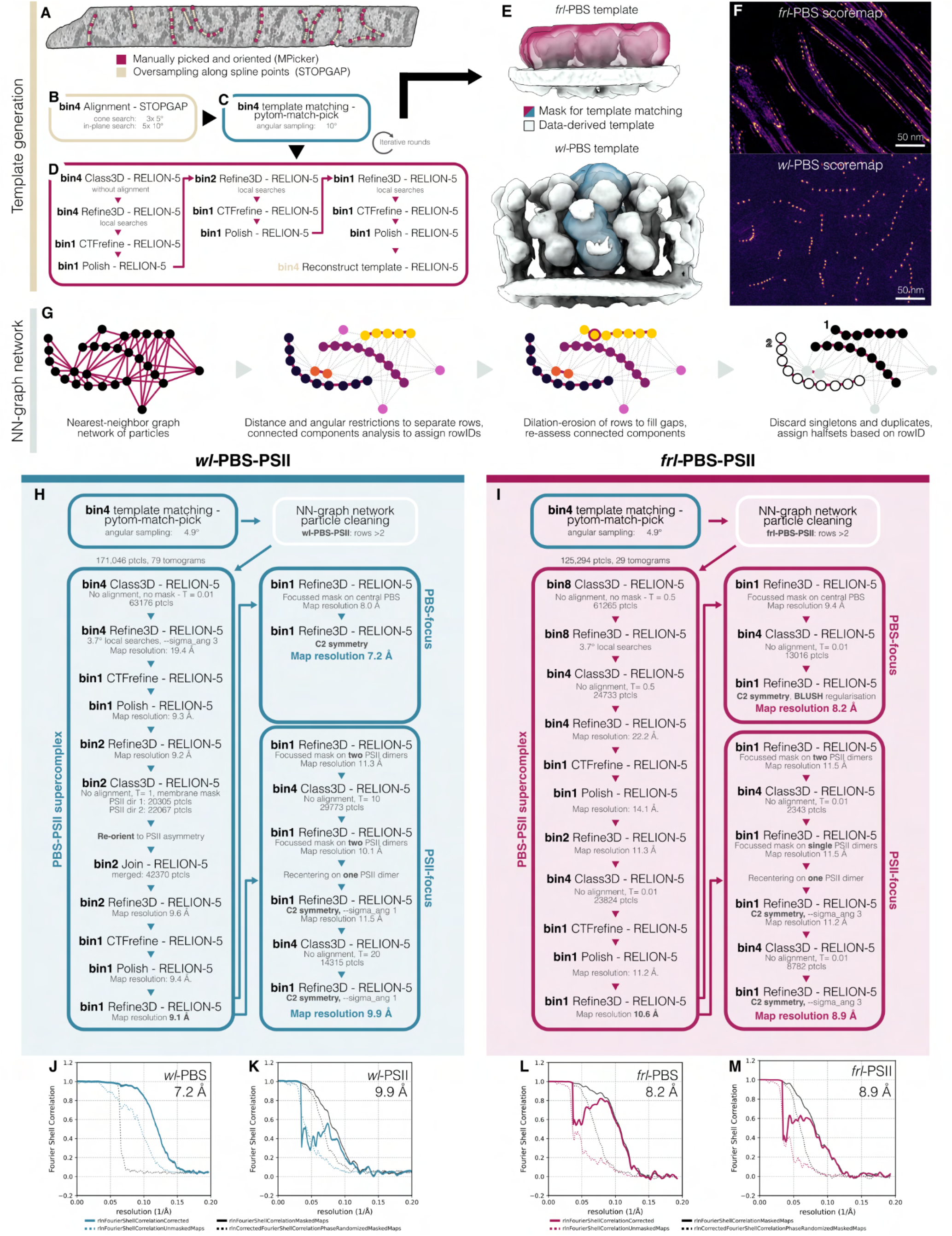
Particle picking and subtomogram averaging workflows for PBS-PSII supercomplexes. **A-F**, Generation of data-derived template for template matching. Points were picked along PBS rows, and splines were oversampled to generate an initial particle set (**A**). These positions were aligned in STOPGAP (**B**), and used as a starting point for template matching with pytom-match-pick (**C**). Particles from initial template matching were then aligned and averaged within RELION5 (**D**). This workflow was iterated until a high-quality template was distilled (**E**). Full row-structures containing three units of PBS-PSII were used for template matching, centered on the central PBS. For FRL data, the central three PBS were masked, shown in red (**E**), while in WL data on the central PBS was masked, shown in blue (**E**). This resulted in score maps with distinct peaks in clear row formations, as expected from the data (**F**). **G**, Particles were subjected to a nearest-neighbor graph network-based cleaning. In this approach, particles in the same row were identified by distance and angular constraints and then separated with connected components analysis. A round of dilation and erosion of row positions was used to fill gaps in rows. Finally, single, unconnected particles, and duplicates were discarded, and rows were assigned half-sets to ensure independent FSC measurements during alignment. **H**, Workflow for subtomogram averaging of *wl-*PBS-PSII in RELION5, see Methods for details. **I**, Workflow for subtomogram averaging of *frl-*PBS-PSII in RELION5, see Methods for details. **J**, Gold standard FSC for focused refinement of *wl-*PBS, final resolution at FSC = 0.143 is 7.2 Å. **K**, Gold standard FSC for focused refinement of *wl-*PSII, final resolution at FSC = 0.143 is 9.9 Å. **L**, Gold standard FSC for focused refinement of *frl-*PBS, final resolution at FSC = 0.143 is 8.2 Å. **M**, Gold standard FSC for focused refinement of *frl-*PSII, final resolution at FSC = 0.143 is 8.9 Å.

**Figure S8.**
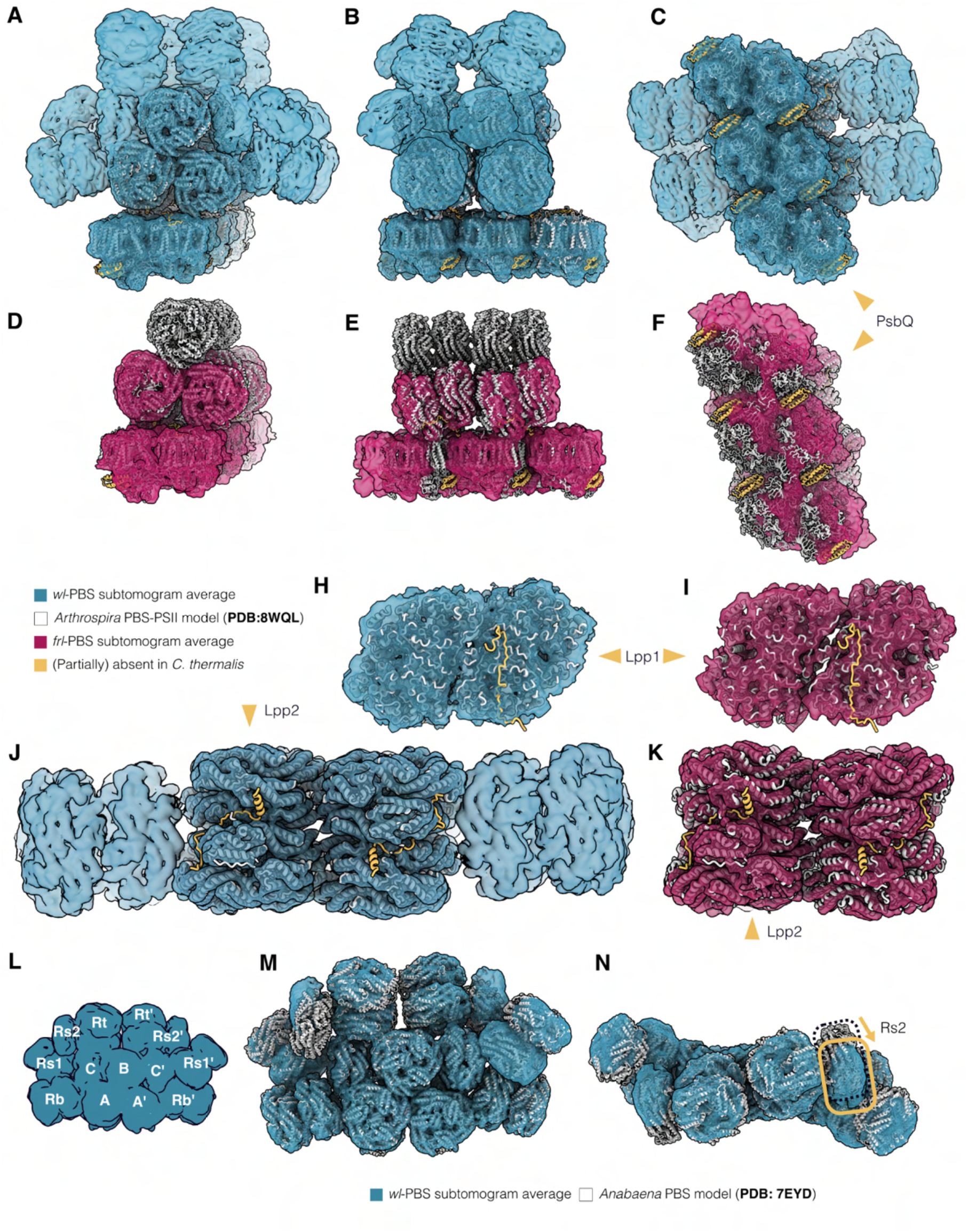
Structural comparison of native *wl-*PBS-PSII and *frl-*PBS-PSII to published structures. **A-C**, Comparison of our *in situ wl-*PBS-PSII supercomplex structure (transparent blue), with the *in situ* PBS-PSII supercomplex structure from *Arthrospira* sp. FACHB-439 (PDB: 8WQL^1^) as seen from front (**A**), side (**B**), and bottom (**C**) views. Notably, PsbQ (in yellow) shows weak density, suggesting partial occupancy, though no distinct classes could be separated. **D-F**, Comparison of our *in situ frl-*PBS-PSII supercomplex structure (transparent red), with the *in situ* PBS-PSII supercomplex structure from *Arthrospira* (PDB: 8WQL^1^) as seen from front (**A**), side (**B**), and bottom (**C**) views. Notably, PBS spacing differs slightly, and PSII organization differs substantially from the model. The PSII subunit PsbQ is completely absent from our *frl-*PSII structure. **H-I**, Rigid body fitting of *wl*-PSII (**H**) and *frl-*PSII (**I**) with PSII from *Arthrospira*. The PSII linker peptide Lpp1 described in *Arthrospira* is absent from the PSII densities in *C. thermalis*. **J-K**, Rigid body fitting of *wl*-PBS (**H**) and *frl-*PBS (**I**) with PBS from *Arthrospira*. The PBS-PSII linker peptide Lpp2 described in *Arthrospira* is absent from the PSII densities in *C. thermalis.* The structures of phycocyanin rods were excluded from the comparison because positioning of the rods differs between the pentacylindrical (*wl-*PBS) and tricylindrical (*Arthrospira*) APC cores. **L**, Description of PBS nomenclature for core and rod complexes based on naming from Zheng et al.^2^ **M-N**, Rigid body fit of cryoEM-based model of the pentacylindrical PBS from *Anabaena* sp. PCC 7120 (PDB: 7EYD^2^) into our *wl-*PBS density, front view (**M**) and top view (**N**). The *in situ* structure of *wl-*PBS is mostly identical to the structure from isolated proteins, apart from a minor repositioning of Rs2 and Rs2’, annotated in yellow.

**Figure S9.**
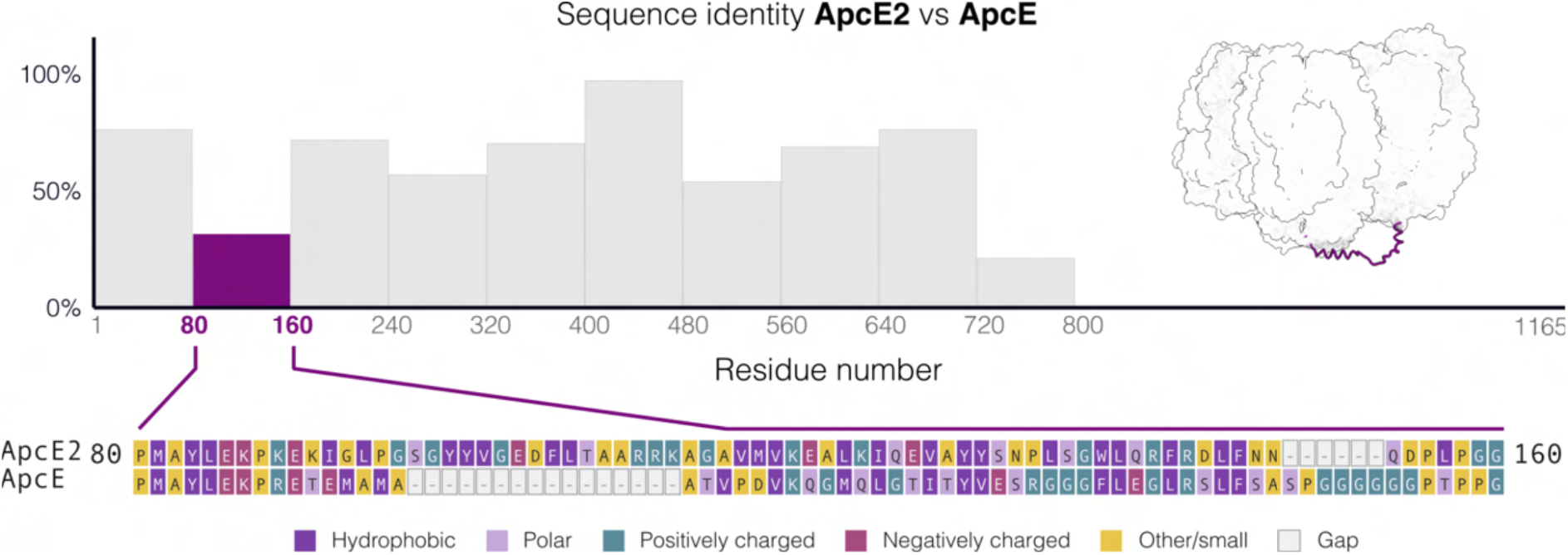
The PSII interaction loop of far-red light ApcE2 is divergent from white light ApcE. Sequence identity between ApcE2 and ApcE across the sequence length. A region of low similarity corresponds to the interaction site with PBS, supporting the role of this region in the differential interaction with PSII between *wl-,* and *frl-*PBS-PSII supercomplexes. The sequence alignment for residues 80-160 (numbered by ApcE2) is shown, and the location of this peptide stretch is depicted in the *frl-*PBS structure.

**Figure S10.**
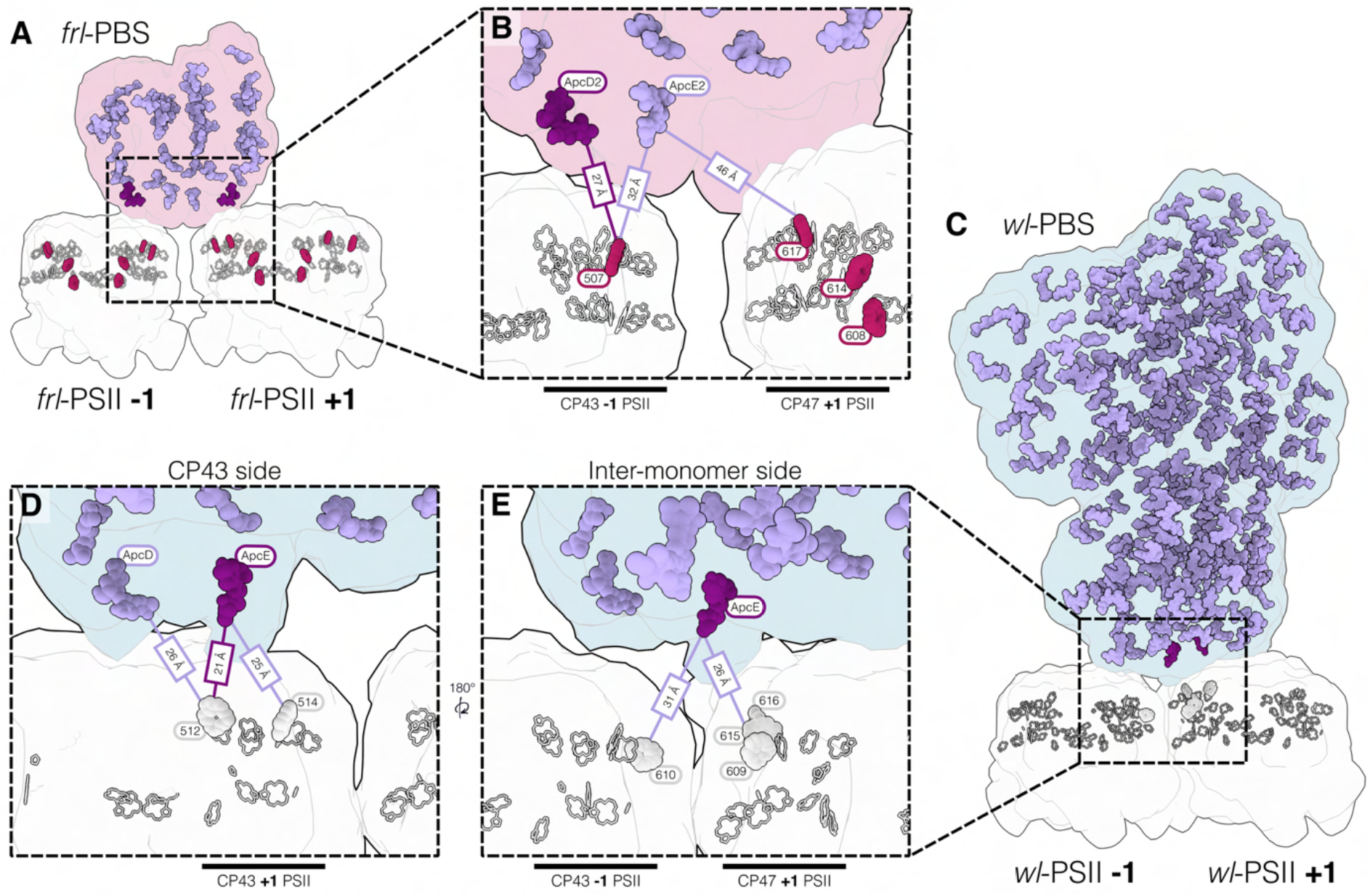
Bilin to chlorophyll distances in *frl-*PBS and *wl-*PBS. **A**, Side view of the *frl*-PBS-PSII supercomplex with the STA density shown as a silhouette. Positions of bilins (purple) and chlorophylls (red and white) in the *frl-*PBS-PSII supercomplex are shown. The closest bilins to Chl *f* are marked in dark purple. Pigment positions are taken from rigid body fitting of PDB: 9I1R^3^ and PDB: 9T5T^3^ into the supercomplex density map. **B**, Zoom into the interaction interface of *frl-*PBS and *frl-*PSII. The closest bilin-Chl *f* positions in -1 and +1 PSII are shown. The shortest distance is found between ApcD2 and Chl *f* 507 (CP43). See **Table S1** for a complete list of pigment distances in *frl-*PBS-PSII. **C**, Side view of the *wl*-PBS-PSII supercomplex with the STA density shown as a silhouette. Positions of bilins (purple) and chlorophylls (white) in the *wl-*PBS-PSII supercomplex are shown. The closest bilins to any of Chl *a* are marked in dark purple. Pigment positions are taken from rigid body fitting of PDB: 7EYD^2^ and PDB: 8IR7^4^ into the supercomplex density map. **D-E**, Zoom into the interaction interface of *wl-*PBS and *wl-*PSII on the CP43 side (**D**) and inter-monomer side (**E**) of the asymmetric interface. The closest bilin-Chl *a* positions in -1 and +1 PSII are shown, and the closest Chl *a* are shown in thicker. The shortest distance is found between ApcE and Chl *a* 512 (CP43) on the CP43 side, and ApcE and Chl *a* 615 on the inter-monomer side. See **Table S2** for a complete list of pigment distances.

**Figure S11.**
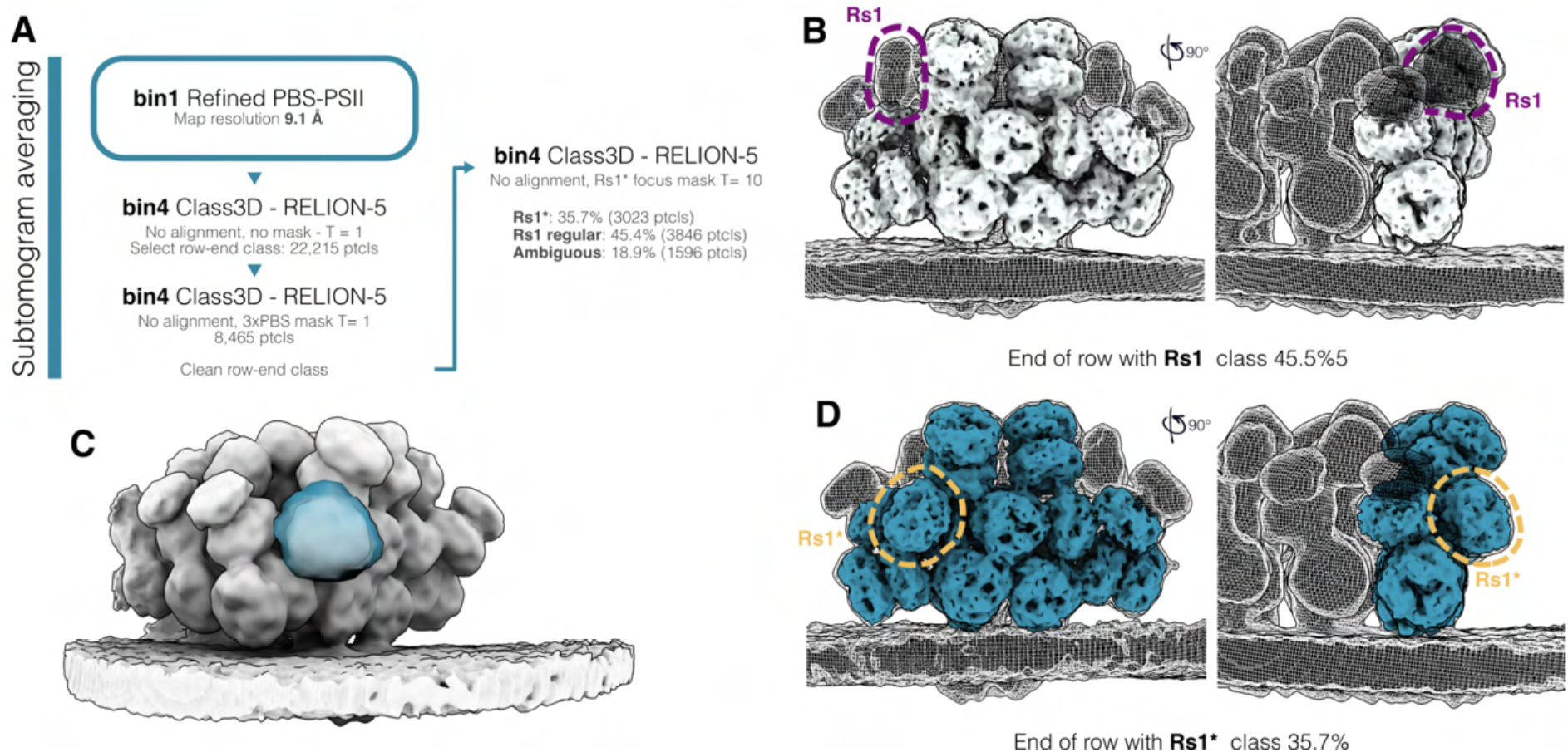
*wl*-PBS row ends show variable organization of the Rs1 rod. **A**, Classification pipeline for the identification of phycobilisome row-ends. **B**, Class representing 46% of row-end particles, showing Rs1 subunit in the conventional position (highlighted in purple). The strongest density is shown as a solid map, lower threshold is shown as a mesh. **C**, Focused mask used in 3D classification. Row end average shown in white, mask shown in blue. **B**, Class representing 36% of row-end particles, showing Rs1 subunit in the lowered Rs1* position (highlighted in yellow). The strongest density is shown as a solid blue map, lower threshold is shown as a mesh.

**Figure S12.**
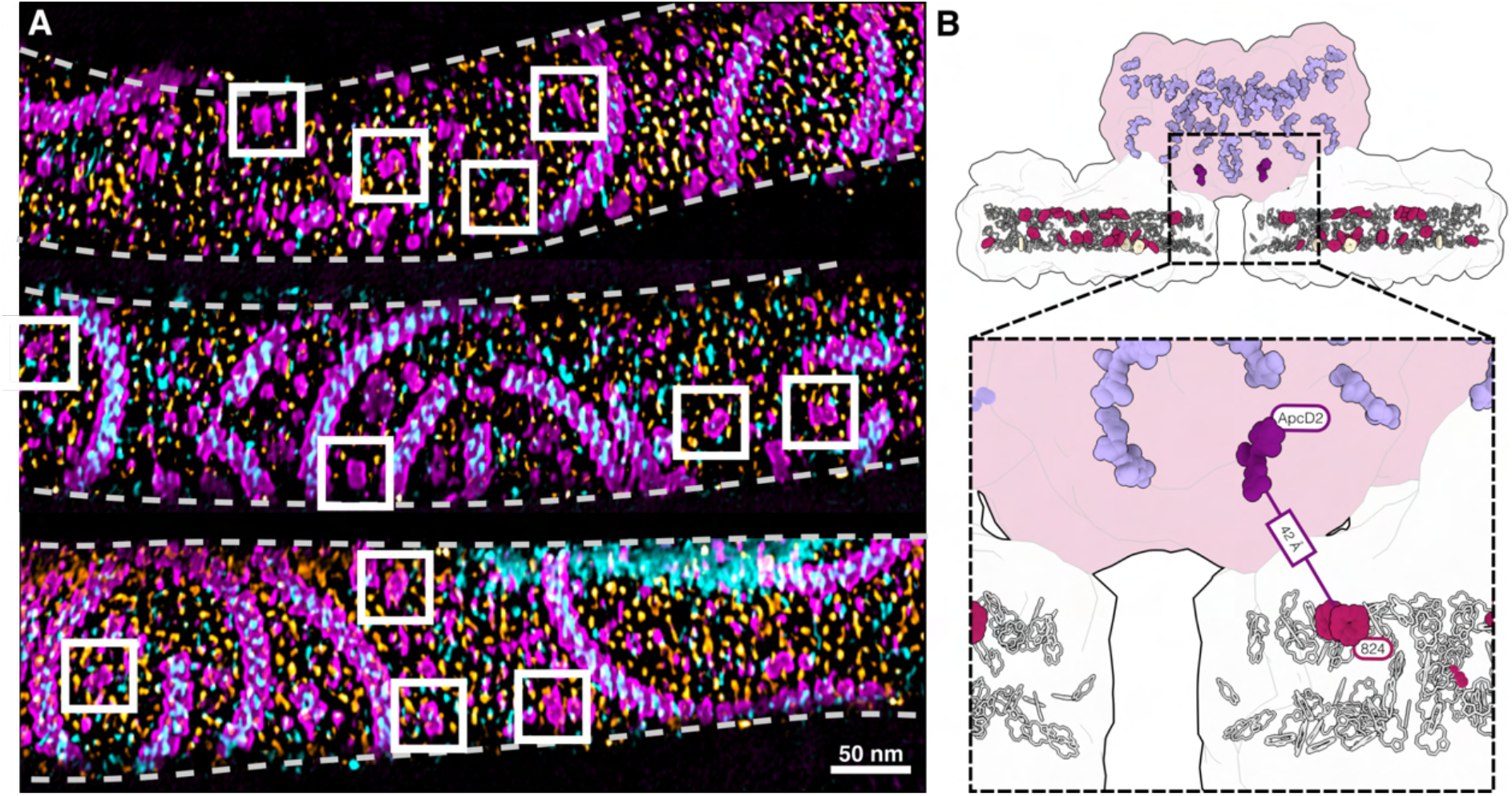
Solitary PBS in FRL forms a *frl-*PBS-PSI supercomplex. **A**, Three examples of membrane surface view overlays with membrane-proximal densities colored as in Figure 4: Densities at ∼6 nm from the thylakoid surface, corresponding to *frl*-PBS, are shown in magenta, while densities at ∼2 nm from the membrane surface, corresponding to PSI, are shown in orange. Densities corresponding to luminal PSII protrusions are in cyan. White boxes highlight examples of solitary *frl*-PBS. Dashed lines mark the edges of the membranes in the image. **B**, Side view of the *frl*-PBS-PSI supercomplex with two trimeric PSI complexes visualized for clarity. STA map density as a silhouette, individual pigments shown in colors: PBS bilins in lavender, Chl *f* in red, Chl *a* in white, and terminal ApcD2 bilins in dark violet. The dashed box highlights the distance between the closest bilin in ApcD2 and the closest Chl *f* (824) of the PSI complex. Pigments positions are taken from rigid body fitting of PDB structures 9I1R^3^ and 9EYS^5^. See **Table S3** for a full list of pigment distances in *frl-*PBS-PSI.

**Figure S13.**
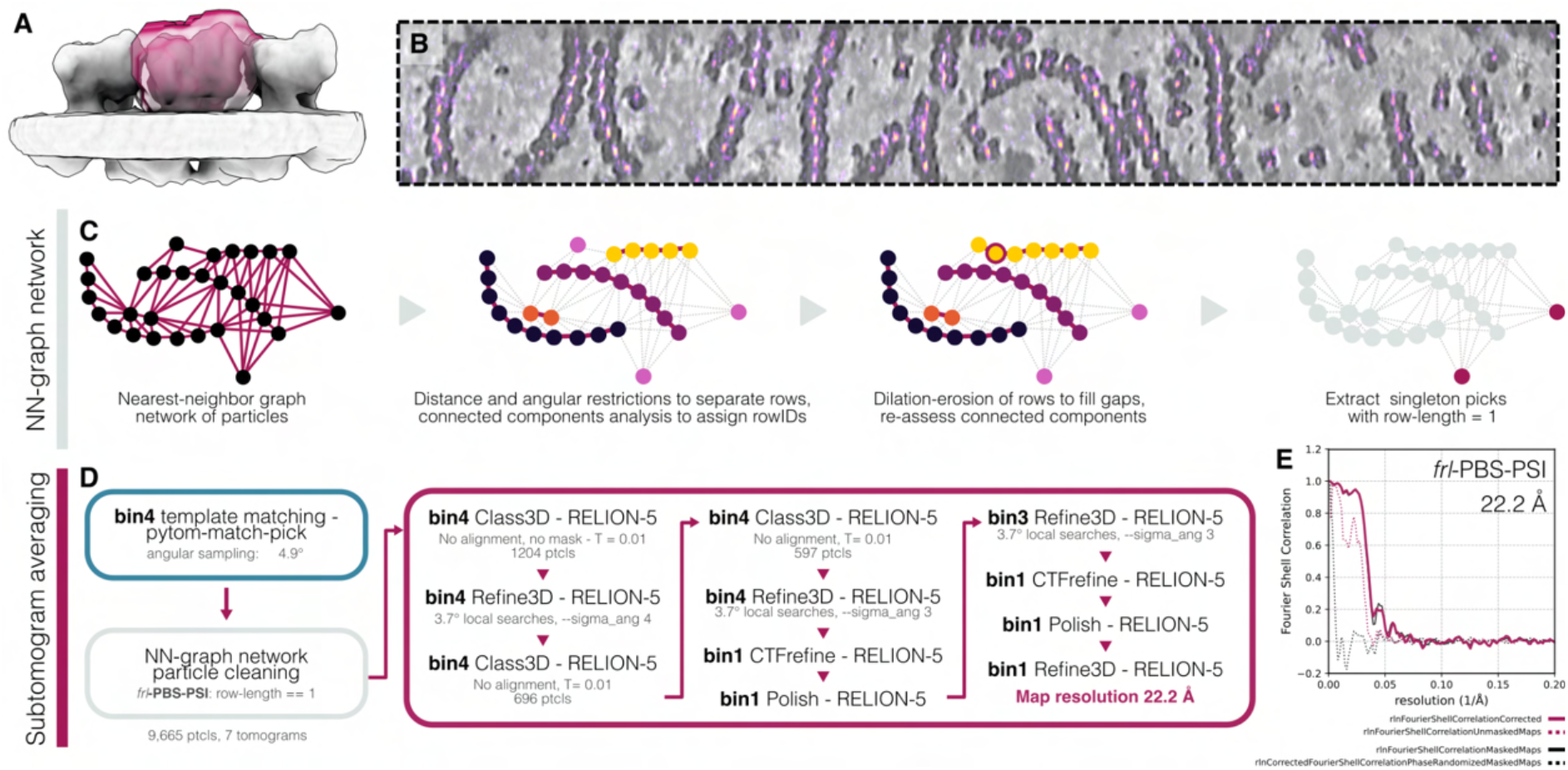
STA processing pipeline for the *frl-*PBS-PSI supercomplex. **A**, *frl-*PBS-PSII was used as a template to identify PBS by template matching. A mask focused on a single PBS (shown in red) was used to ensure picking of solitary PBS. **B**, Overlay of cross-correlation scores from template matching on a flattened membrane surface view, showing clean peaks over PBS, including solitary PBS. Tomogram with PBS densities shown in grayscale, cross correlation scores shown in magenta-to-yellow *Inferno* colormap. **C**, Description of nearest neighbor graph network workflow to identify solitary PBS. Similar to the *frl-*PBS-PSII pipeline, nearest neighbor analysis with a dilation and erosion step was applied to group particles into their rows. Particles with a row length of 1 were extracted as single, unconnected, picks. **D**, Subtomogram averaging and alignment pipeline in RELION5 for the PBS-PSI supercomplex. **E**, Gold standard FSC plot of the *frl*-PBS-PSI average, final global resolution at FSC = 0.143 is 22.2 Å.

**Figure S14.**
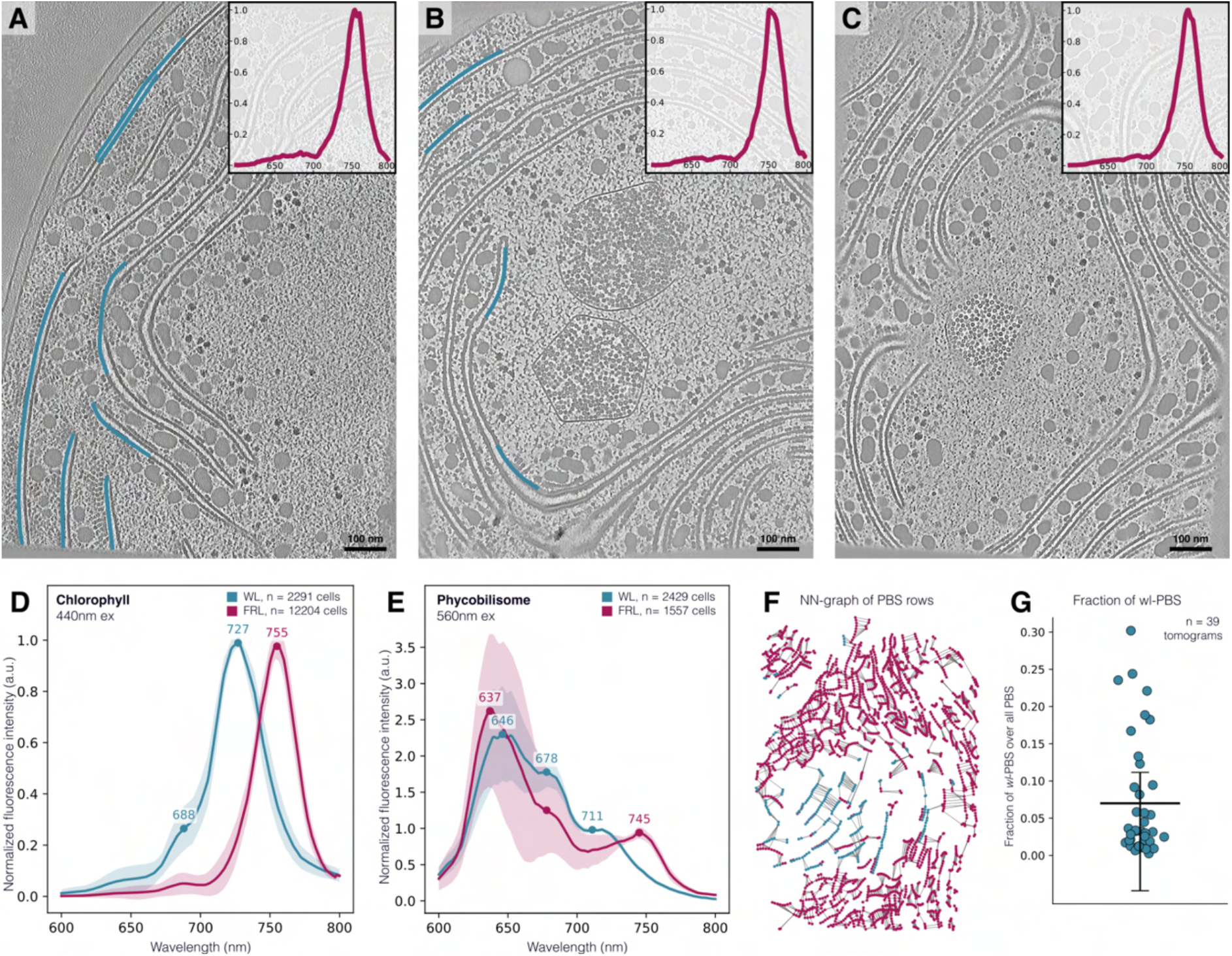
hsCLEM shows *wl-*PBS are present in *C. thermalis* cells with fully red-shifted fluorescence signatures. Three additional examples of tomograms with single cell spectrum, range of abundance of wl-PBS, global hsCLEM spectra, nearest-neighbor (NN) graph visualization for *wl*-rows segregation, and fraction of *wl*-PBS scatterplot. **A, B, C**, Cryo-ET slices through three tomograms of cells acclimated to FRL. Respective insets show corresponding single cell spectra obtained from hsCLEM. Blue membrane highlights in (**A**) and (**B**) point to the *wl*-PBS present in the cells. **D**, hsCLEM fluorescence spectra using chlorophyll excitation at 440nm from WL- (blue, n=2291) and FRL-acclimated cells (red, n=12204). Maximal emission at 688 nm (*wl-*PSII), and 727 nm (*wl-*PSI), and 755 nm (*frl*) are highlighted on the plot. **E**, hsCLEM fluorescence spectra using phycobilisome excitation at 560nm from WL- (n=2429, blue) and FRL-acclimated cells (n=1557, red). Respective local emission maxima are highlighted on the plot. The terminal emission for FRL-grown cells is clearly shifted to longer wavelengths. **F**, Visualization of the NN-distribution of PBS rows in a tomogram from the FRL-acclimated cells that retained some *wl*-PBS. Positions of *frl*-PBS in red, *wl*-PBS in blue. **G**, Plot showing fraction of *wl*-PBS (mean: 0.07, median 0.03) in the tomograms from the FRL-acclimated cells from n=39 tomograms. Each point represents the fraction of *wl-*PBS over total PBS content in a single tomogram. Mean and standard deviation are displayed.

**Figure S15.**
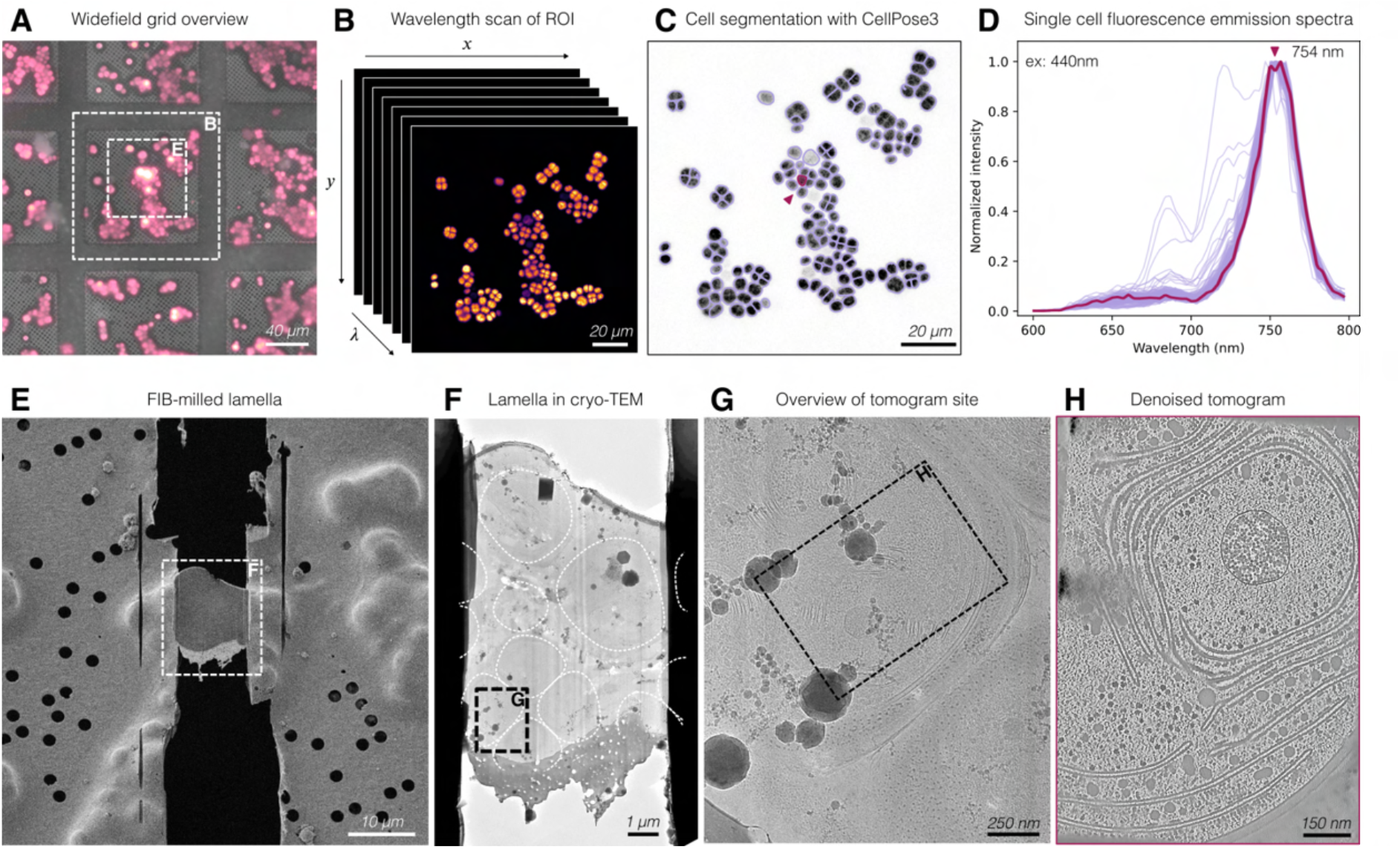
Extended workflow for acquisition and analysis of hsCLEM data. **A**, Wide field overview of *C. thermalis* cells on a grid in a cryo fluorescence microscope. Reflective light image shown in grayscale, autofluorescence shown in magenta-to-yellow intensity (Inferno) colormap (ex: 541-551 nm, em: 565-605 nm). Scale bar 40 µm. **B**, Representation of a wavelength scan collected with a cryo confocal microscope. Images are collected from 600-800 nm with a 3 nm stepsize and 15 nm bandwidth. Scale bar 20 µm. **C**, Maximum intensity projection of the wavelength scan is used to segment individual cells using CellPose3^2^. Fluorescence is shown in greyscale, segmented outlines are shown in lilac. Each cell is assigned a unique cell ID, which can be used to link individual cell spectra to the tomographic data. An example cell is highlighted in red, and corresponds to the red spectrum in (**D**) and tomogram shown in (**H**). Scale bar 20 µm. **D**, Single cell fluorescence emission traces from wavelength scan shown in (**C**). The highlighted cell in (**C**) is shown in red. Single cell traces reveal there is slight heterogeneity in the sample. **E**, Scanning electron microscopy (SEM) image of a thinned lamella at the position denoted in (**A**). Scale bar 10 µm. **F**, Image of lamella from (**E**) in a transmission electron microscope (TEM) prior to tilt series acquisition. Cell outlines from the fluorescence cell segmentation in (**C**) are shown as dashed white lines. Scale bar 1 µm. **G**, Zoom-in on the lamella overview from (**F**) showing the site of tomogram acquisition (black dashed box). Scale bar 250 nm. **H**, Tomogram collected at the site marked in (**G**), corresponding to the highlighted cell and spectrum in (**C**) and (**D**). Scale bar 150nm.

**Figure S16.**
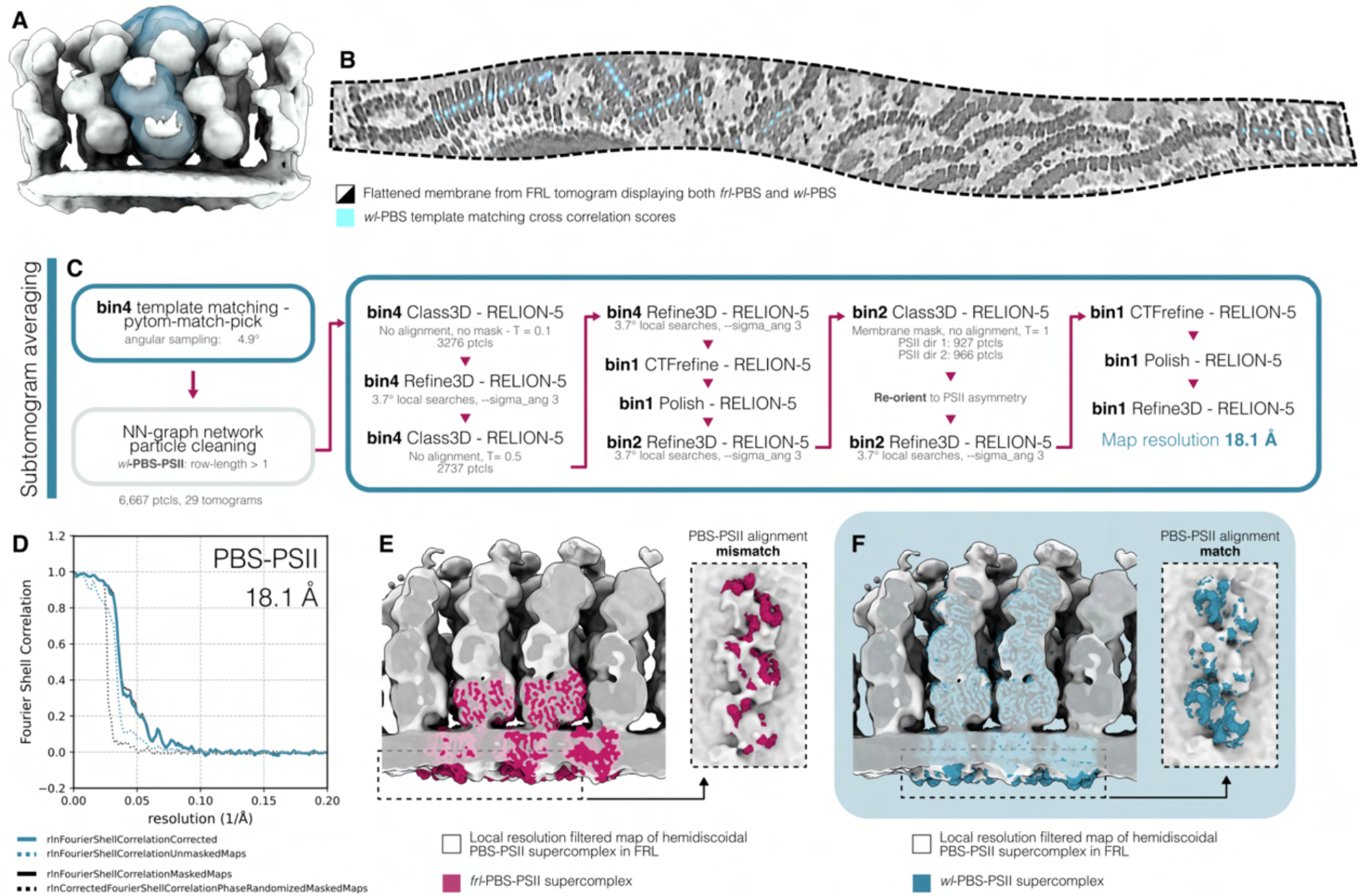
Subtomogram averaging workflow for hemidiscoidal PBS-PSII found in FRL. **A**, *wl-*PBS template was used for template matching of hemidiscoidal PBS-PSII in FRL. A mask was focused on the central PBS and did not include PSII density to avoid bias towards row organization. **B**, Overlay of cross-correlation scores from template matching on a flattened membrane surface view. Tomogram with *wl-*PBS and *frl-*PBS densities shown in grayscale, template matching cross-correlation scores shown in cyan. Despite shared membrane domains, template matching was highly specific for hemidiscoidal PBS structures. **C**, Processing pipeline for the subtomogram averaging and alignment in RELION5 of *wl-*PBS-PSII from FRL tomograms. **D**, Gold standard FSC curves for *wl-*PBS-PSII supercomplex in FRL. Final global resolution at FSC = 0.143 is 18.8Å. **E-F**, Hemidiscoidal PBS-PSII (white) structure from FRL cells compared to *frl-*PBS-PSII (red) and *wl-*PBS-PSII (blue) complexes. The bottom luminal surface view of the membrane (insets) is shown to display the difference in PSII positioning. While *frl-*PBS-PSII shows a mismatch, *wl-*PBS-PSII is in full agreement with the organization of the PBS-PSII supercomplex in FRL conditions.

## Supplementary tables

**Table S1.** Donor-acceptor pigment distances in *frl*-PBS-PSII. Distances calculated from rigid body docking of *frl-*PSII (PDB: 9T5T) and *frl-*PBS (PDB: 9I1R) into local resolution filtered supercomplex map. Residue numbers are as pigment annotations used in PDBs. Chl *f* acceptor sites are highlighted in red, CP43 sites are shown in white, and CP47 sites are shown in grey.

| Donor protein | Donor residue | Acceptor protein | Acceptor pigment | Acceptor residue | Edge distance (Å) | Centroid distance (Å) |
| --- | --- | --- | --- | --- | --- | --- |
| ApcD3 | 201 | PsbC2 (CP43) | CLA | 511 | 18.7 | 30.8 |
| ApcE2 | 801 | PsbC2 (CP43) | CLA | 513 | 21.4 | 34.5 |
| ApcD3 | 201 | PsbC2 (CP43) | CLA | 509 | 24.2 | 35.7 |
| ApcD3 | 201 | PsbC2 (CP43) | CLA | 508 | 24.6 | 34.4 |
| ApcE2 | 801 | PsbC2 (CP43) | CLA | 512 | 24.8 | 39.3 |
| ApcD3 | 201 | PsbC2 (CP43) | CLA | 512 | 25.2 | 38.7 |
| ApcD3 | 201 | PsbC2 (CP43) | CLA | 513 | 25.8 | 36.1 |
| ApcD3 | 201 | PsbC2 (CP43) | F6C | 507 | 27.4 | 39.5 |
| ApcD3 | 201 | PsbC2 (CP43) | CLA | 510 | 28.8 | 41.0 |
| ApcD3 | 201 | PsbC2 (CP43) | CLA | 506 | 30.6 | 43.1 |
| ApcD3 | 201 | PsbC2 (CP43) | CLA | 505 | 30.7 | 41.7 |
| ApcE2 | 801 | PsbC2 (CP43) | F6C | 507 | 32.0 | 44.8 |
| ApcE2 | 801 | PsbC2 (CP43) | CLA | 511 | 32.7 | 44.1 |
| ApcE2 | 801 | PsbC2 (CP43) | CLA | 509 | 33.9 | 44.4 |
| ApcB2 | 201 | PsbC2 (CP43) | CLA | 511 | 34.2 | 46.8 |
| ApcE2 | 801 | PsbC2 (CP43) | CLA | 506 | 34.6 | 44.9 |
| ApcE2 | 801 | PsbC2 (CP43) | CLA | 508 | 37.8 | 49.2 |
| ApcD5 | 201 | PsbB2 (CP47) | CLA | 610 | 38.2 | 49.3 |
| ApcD5 | 201 | PsbB2 (CP47) | CLA | 611 | 39.3 | 51.9 |
| ApcD3 | 201 | PsbC2 (CP43) | CLA | 503 | 39.8 | 52.2 |
| ApcD5 | 201 | PsbB2 (CP47) | CLA | 616 | 39.8 | 51.4 |
| ApcE2 | 801 | PsbC2 (CP43) | CLA | 503 | 40.0 | 54.5 |

**Table S2.**
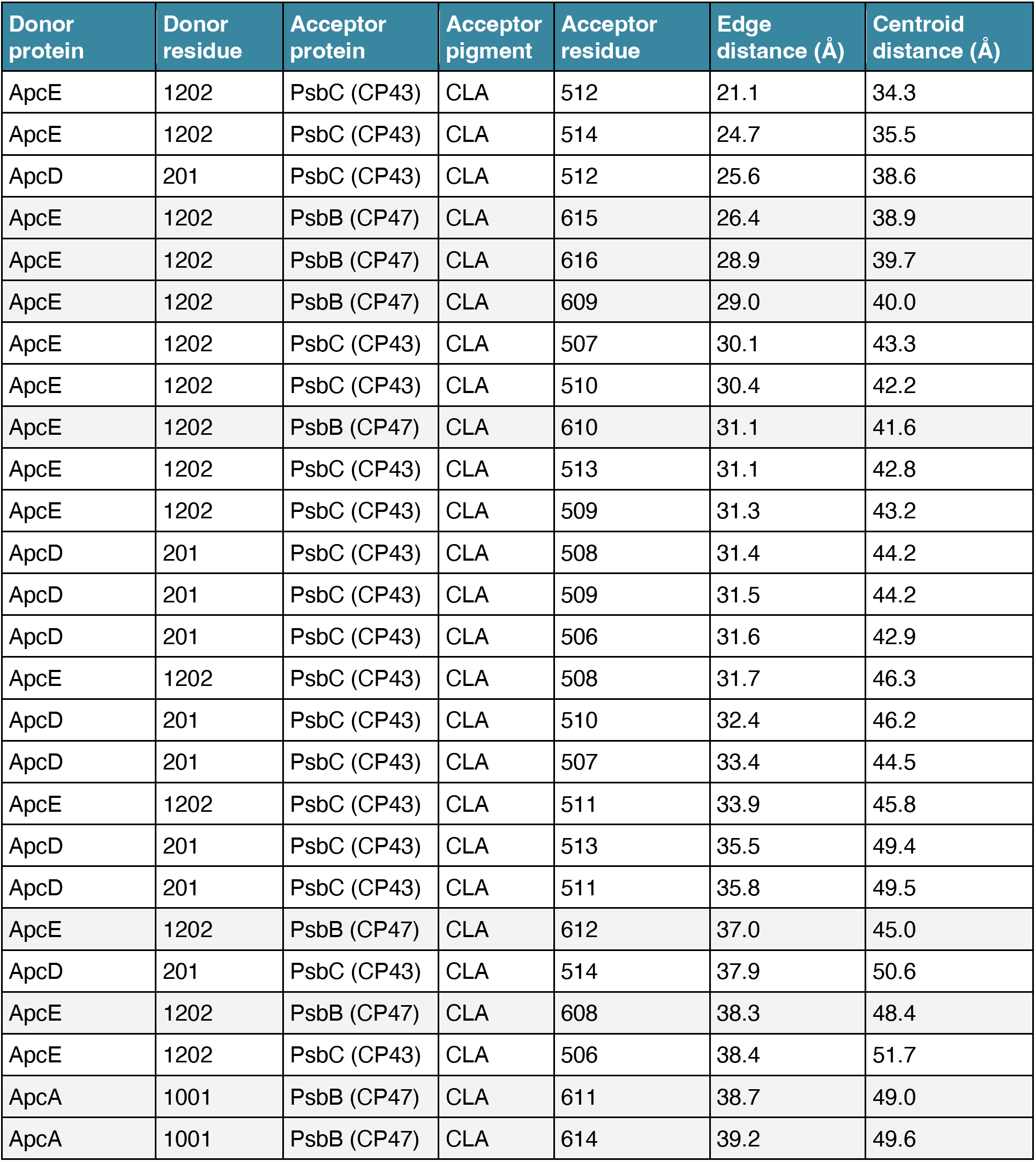
Donor-acceptor pigment distances in *wl*-PBS-PSII. Distances calculated from rigid body docking of PSII (PDB: 7N8O) and pentacylindrical PBS (PDB: 7EYD) into local resolution filtered supercomplex map. Residue numbers are as pigment annotations used in PDBs. CP43 acceptor sites are shown in white and CP47 sites are shown in grey.

**Table S3.** Donor-acceptor pigment distances in *frl*-PBS-PSI. Distances calculated from rigid body docking of *frl-*PSI (PDB: 9EYS) and *frl-*PBS (PDB: 9I1R) into local resolution filtered supercomplex map. Residue numbers are as pigment annotations used in PDBs. Chl *f* sites are highlighted in red, CP43 sites are shown in white and CP47 sites are shown in grey.

| Donor protein | Donor residue | Acceptor protein | Acceptor pigment | Acceptor residue | Edge distance (Å) | Centroid distance (Å) |
| --- | --- | --- | --- | --- | --- | --- |
| ApcE2 | 801 | PsaA2 | CLA | 821 | 25.0 | 38.1 |
| ApcE2 | 801 | PsaA2 | CLA | 822 | 26.8 | 39.9 |
| ApcE2 | 801 | PsaA2 | CLA | 820 | 27.4 | 40.3 |
| ApcE2 | 801 | PsaA2 | CLA | 812 | 28.5 | 39.1 |
| ApcE2 | 801 | PsaA2 | CLA | 824 | 31.3 | 43.7 |
| ApcE2 | 801 | PsaA2 | CLA | 813 | 32.4 | 45.1 |
| ApcE2 | 801 | PsaA2 | CLA | 811 | 32.9 | 43.5 |
| ApcE2 | 801 | PsaA2 | CLA | 823 | 33.0 | 44.2 |
| ApcE2 | 801 | PsaA2 | CLA | 819 | 33.0 | 44.7 |
| ApcD3 | 201 | PsaB2 | CLA | 812 | 37.4 | 44.0 |
| ApcD3 | 201 | PsaA2 | CLA | 821 | 37.6 | 49.0 |
| ApcD5 (X/x) | 201 | PsaA2 | CLA | 811 | 37.8 | 47.5 |
| ApcE2 | 801 | PsaB2 | CLA | 823 | 38.5 | 49.8 |
| ApcD3 | 201 | PsaB2 | CLA | 811 | 39.1 | 49.2 |
| ApcD5 (X/x) | 201 | PsaB2 | CLA | 832 | 39.3 | 52.3 |
| ApcD5 (X/x) | 201 | PsaA2 | CLA | 812 | 40.6 | 50.0 |
| ApcD3 | 201 | PsaA2 | CLA | 822 | 40.7 | 51.4 |
| ApcE2 | 801 | PsaA2 | CLA | 830 | 40.8 | 52.2 |
| ApcE2 | 801 | PsaA2 | CLA | 805 | 41.0 | 50.0 |
| ApcD5 (X/x) | 201 | PsaA2 | CLA | 804 | 41.5 | 53.2 |
| ApcD5 (O/o) | 201 | PsaA2 | CLA | 804 | 41.6 | 55.3 |
| ApcD3 | 201 | PsaA2 | CLA | 820 | 41.6 | 54.5 |
| ApcE2 | 801 | PsaB2 | F6C | 824 | 41.8 | 51.6 |
| ApcD5 (X/x) | 201 | PsaB2 | F6C | 824 | 42.0 | 51.6 |
| ApcD5 (X/x) | 201 | PsaA2 | CLA | 805 | 42.6 | 53.3 |
| ApcD5 (X/x) | 201 | PsaA2 | CLA | 813 | 42.7 | 52.3 |
| ApcD3 | 201 | PsaB2 | CLA | 805 | 43.0 | 52.0 |
| ApcD5 (X/x) | 201 | PsaA2 | CLA | 824 | 43.2 | 54.9 |
| ApcD5 (O/o) | 201 | PsaB2 | CLA | 832 | 43.2 | 54.9 |
| ApcD5 (X/x) | 201 | PsaB2 | CLA | 833 | 43.7 | 57.4 |
| ApcD5 (O/o) | 201 | PsaB2 | CLA | 833 | 43.7 | 55.1 |

